# Glypican-6 deficiency causes dose-dependent conotruncal congenital heart malformations through abnormal remodelling of the endocardial cushions

**DOI:** 10.1101/2021.06.28.450191

**Authors:** Gennadiy Tenin, Alexander Crozier, Kathryn E. Hentges, Bernard Keavney

**Author notes:** Division of Biosciences, University College London, London, UK. Address correspondence to: Gennadiy Tenin, Division of Cardiovascular Sciences, Faculty of Biology, Medicine and Health, University of Manchester, Upper Brook Street, Manchester, UK, M13 9PT.

## Abstract

Tetralogy of Fallot (TOF) is considered to be the commonest type of cyanotic congenital heart disease (CHD). A previous GWAS showed significant association between TOF and single nucleotide polymorphisms in chromosome 13q31. Here through integration of population genomic and chromosomal interaction data we identify the heparan sulfate proteoglycan glypican-6 (GPC6) as the potentially responsible gene at the associated locus. We showed that GPC6 is expressed in the endocardial cushions at the appropriate time in development to contribute to TOF risk. We generated mice homozygous for a Gpc6 KO allele, which exhibit 100% neonatal lethality with severe cardiac malformations, namely TOF-type double outlet right ventricle (DORV) with rightward mal-positioned aorta and perimembranous ventricular septal defect (VSD), together with right ventricular (RV) hypertrophy and narrowing of the pulmonary artery. We established a dose-response relationship between Gpc6 expression and the anatomical severity of cardiac malformations. We showed the mouse knockout phenotype arises from abnormal morphology of the endocardial cushions, and tissue-specific knockout of Gpc6 in endothelial and neural crest cell lineages produces a phenotype featuring VSD and aortic malposition analogous to human TOF. This successful identification of a CHD gene from GWAS data suggests that larger GWA studies may find additional causative genes.

## Introduction

Congenital heart defects (CHD) are the commonest anomalies present at birth and affect around 1% of new-borns (1). Tetralogy of Fallot (TOF) is considered to be the commonest type of cyanotic congenital heart disease with an incidence of 2-3 per 10,000 live births (2). TOF consists of four anatomical abnormalities of the heart: ventricular septal defect (VSD), overriding aorta (OAo), pulmonic stenosis (PS) and right ventricular hypertrophy (2). Although surgical treatment of TOF yields adult survival in over 90% of patients, it is still accompanied by late morbidity and mortality, including sudden cardiac death (3, 4). A number of studies have identified both environmental (e.g. diabetes mellitus (5)) and genetic factors involved in the development of TOF; the latter include chromosomal abnormalities such as 22q11 deletion syndrome, as well as rare variants in genes such as *NKX2.5, TBX1, GATA4, and Jagged1*, that are involved in the proliferation, differentiation, and migration of cardiac cells during development (6–12). However, these only explain a minority of TOF; in the approximately 80% of cases that are non-syndromic and sporadic, the cause is unknown (13). Familial recurrence risk studies have consistently indicated a genetic predisposition to TOF (14, 15); in keeping with this, studies have shown an excess of rare copy number variants (10, 16–18) and of rare deleterious single-nucleotide variants, in TOF patients (19–24). Regarding common genetic variation, a 2013 genome-wide association study (GWAS) identified 3 regions significantly associated with sporadic, non-syndromic TOF (25). In this study, we investigate one of these loci, chr13q31, which has the highest degree of association (p = 3.03e-11). We identify the nearby gene *GPC6* as a critical gene for cardiac outflow tract development, which is the overwhelmingly likely cause of the GWAS signal.

## Results

### Genomic and epigenetic data in the chr13q31 locus associated with TOF implicates *GPC6*

Only two protein coding genes, *GLYPICAN 5* and *GLYPICAN 6*, are located within 2Mb of the GWAS lead SNP rs7982677 (Figure 1a). A large 3.7 Mb gene desert lies upstream of *GPC5*; downstream of *GPC6*, the next coding gene, *DCT*, is located over 2 Mb away from rs7982677. rs7982677 variant is located in intron 7-8 of *GPC5* and is in high linkage disequilibrium (r^2^>0.8) with 44 other genomic variants extending over a 66 Kb region (Figure 1a). No variant has an effect on the protein sequence of GPC5. Analysis of the GTEx data (gtexportal.org, v05-08-15) did not reveal significant eQTL association between these variants and expression of any gene in any tissue. A human-specific, non-coding RNA, *GPC5-AS2*, lies in the LD locus: we showed that it was not expressed in the human embryonic heart at any stage relevant to the development of TOF (Supplemental Figure 1); it was therefore discarded as a candidate gene.

**Figure 1.**
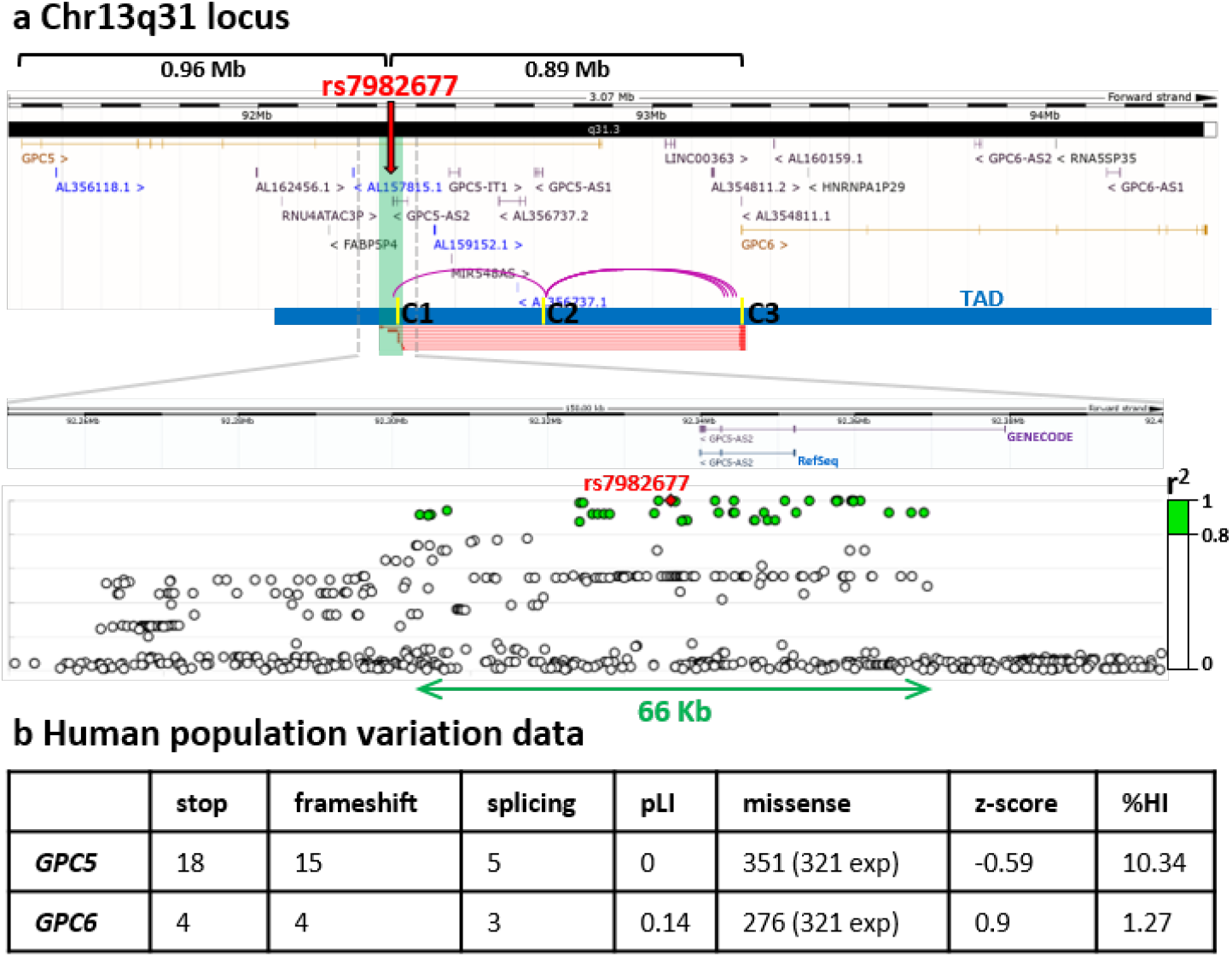
Analysis of the Chr13q31 locus. **a** The overview of the locus (GRCh38 human genome annotation). The distance between rs7982677 and start sites of *GPC5* and *GPC6* is shown in megabases (Mb). Red arrow denotes the location of the rs7982677 variant within the haploblock (green bar). Blue bar - topologically associated domain (TAD). Red lines represent the interaction loops between the GPC6 promoter region and the haploblock. Yellow marks show the location of CTCF binding sites (C1-C3). Violet arcs denote highly correlated DHS sites. Magnified LD region is shown at the bottom, with green circles representing variants in high linkage disequilibrium with rs7982677 (r^2^>0.8). GENECODE and RefSeq annotations of the GPC5-AS2 non-coding RNA are shown. **b** The number of damaging mutations found in *GPC5* and *GPC6* genes (gnomAD 2.1.1), including loss-of-function (stop codons, frameshift mutations and mutations affecting splicing) and missense mutations. pLI is a probability of LOF intolerance (values between 0 (tolerant) and 1 (intolerant)). Z-score describes the degree to which a gene is depleted of missense variants compared to expected values (positive value predicts intolerance to missense mutations). %HI haploinsufficiency score is based on copy-number variant data (a score of 0-10 suggests haploinsufficiency).

We analysed the structure of topologically associated domains (TADs) in chr13q31.3, based on chromosomal interaction HiC data (26), as such domains usually contain *cis*-regulatory sequences of genes within the TAD. In 37 cells/tissues, a TAD has been demonstrated starting 300-500 Kb upstream of rs7982677, and ending downstream of the 3’ terminus of the *GPC6* gene (Supplemental Figure 2, black labels), suggesting a high probability of functional association between the rs7982677 haploblock and *GCP6*, but not *GPC5*. Promoter Capture HiC identified 7 direct chromosomal interactions between the *GPC6* promoter and the haploblock (denoted in red lines in Figure 1a); no similar interactions were found for the *GPC5* promoter. Analysis of open chromatin data (27) identified the presence of three highly correlated (r>0.7) DNase I hypersensitive sites (DHSs), connecting the GWAS locus to the GPC6 promoter region (violet arcs in Figure 1a); no such interactions exist for the GPC5 promoter. ENCODE data shows three CTCF binding sites located in close proximity to the DHSs (yellow marks C1-C3 in Figure 1a) whose occupancy positively correlates with GPC6 gene expression in 13 ENCODE cell lines studied (Pearson correlation coefficient r = 0.5974, 0.6596, 0.8462, p = 0.031083, 0.014174, 0.000266 for C1, C2 and C3, correspondingly).

Population genomic data from gnomAD v2.1.1 (gnomad.broadinstitute.org) (28) shows the Probability of Loss-of-function Intolerance score (pLI) for GPC6 is 0.14 and 0 for GPC5, predicting *GPC6* being more likely loss-of function intolerant than *GPC5* (Figure 1b). In keeping with this, the DECIPHER (decipher.sanger.ac.uk) (29) haploinsufficiency score (%HI) of 1.27 for *GPC6* and 10.34 for *GPC5* suggests that *GPC6*, but not *GPC5*, is likely to be haploinsufficient (Figure 1b). Additionally, based on the Z-score value (gnomAD), *GPC6* is more likely to be intolerant to missense changes than *GPC5* (0.9 vs −0.59, positive value predicts intolerance; Figure 1b).

Lastly, we considered the previously reported data from available mouse knock-out (KO) studies of both genes. A *Gpc5^tm1aWtsi^* KO strain (MGI:4455511) has been produced by WTSI, and homozygous animals were found to be viable with mild abnormalities in rib cage morphology. In contrast, mice homozygous for the *Gpc6^tm1Lex^* KO allele (MGI:5007162) (30) were reported as lethal, although no cardiac abnormalities were specifically described.

The analysis of the genes in the chr13q31.3 locus indicated that the *GPC6* gene was the most plausible candidate gene accounting for the GWAS result.

### Expression analysis of GPC6 in the developing heart

We first analysed the expression of *GPC6/Gpc6* in human and mouse cardiac tissues by PCR. In human heart cDNA, *GPC6* expression was detected at Carnegie stages (CS) 16, 19 and 21, but not CS12 or 23 (Figure 2a). In the mouse hearts, *Gpc6* was found at embryonic days (E) 10.5 to 14.5 but not at E9.5. *Gpc6* was therefore detected at similar stages of heart morphogenesis in mouse and human (color-coded in Figure 2a).

**Figure 2.**
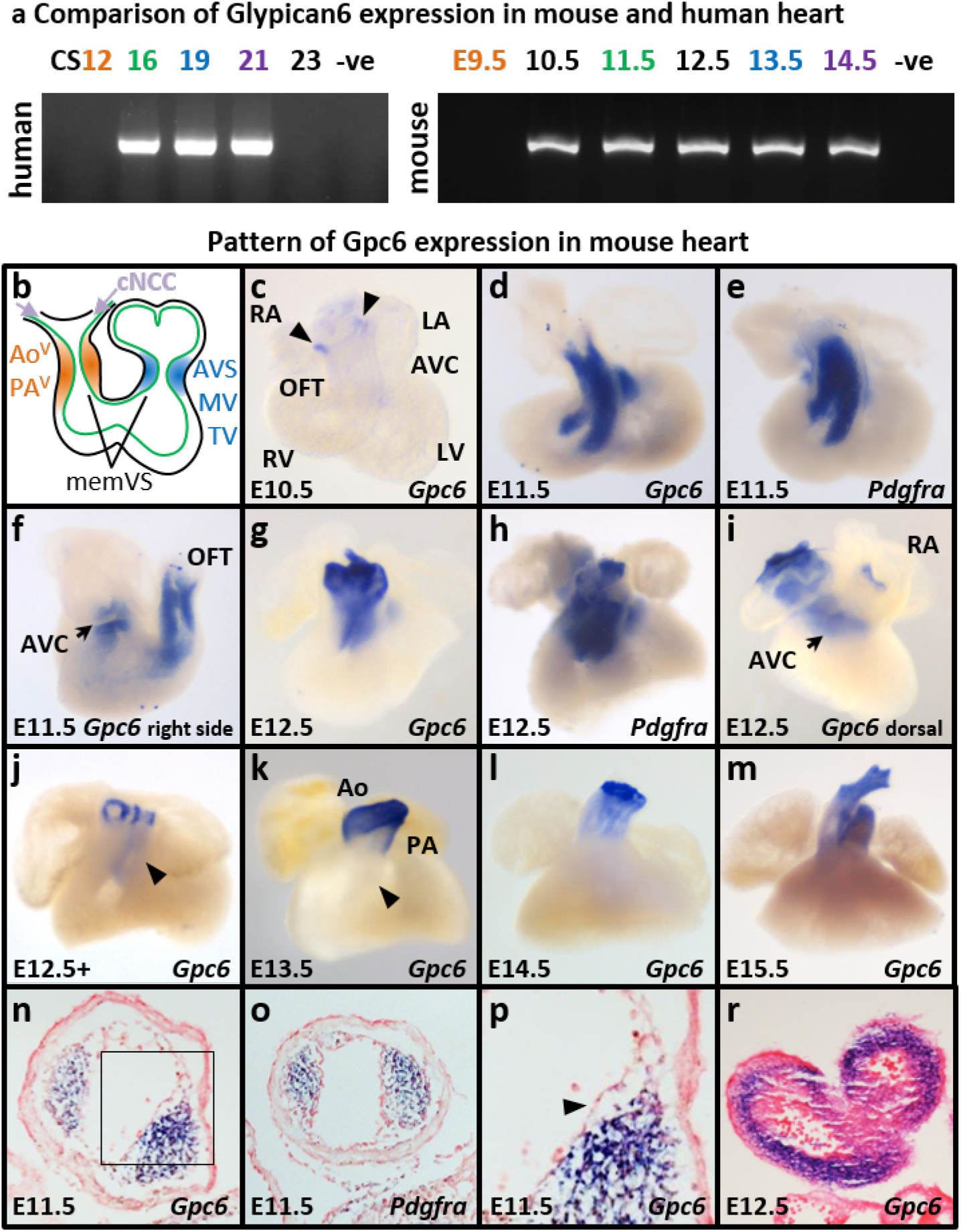
Analysis of GPC6 expression in human and mouse embryonic hearts. **a** Comparison of Glypican 6 expression in human and mouse heart, analysed by PCR. CS Carnegie stages of human embryonic development, E days of mouse embryonic development, -ve negative control. Similar stages of cardiogenesis between two species are indicated with the same colour. **b** Schematics of the mouse embryonic heart. Endocardium is shown in green, OFT cushions in orange, AV cushions in blue. Violet arrows show the direction of cardiac neural crest cells migration (cNCC). memVS membranous ventricular septum, Ao^V^ aortic valve, PA^V^ pulmonary artery valve, AVS atrioventricular septum, MV mital valve, TV tricuspid valve. **c-m** Pattern of gene expression in mouse embryonic heart at E10.5-15.5: *Gpc6* in **c**, **d**, **f**, **g**, **i-m**, *Pdgfra* expression in e, h. RA, LA right/left atrium, RV, LV right/left ventricle, OFT outflow tract, AVC atrioventricular canal. Arrowheads in **c** point to aortic bulb, in **j-k** to conal septum. **n-r** histological sections highlighting the expression of *Gpc6* and *Pdgfra* in the mesenchyme of E11.5 EC (blue in **n-p**) but not in the endocardium (endo) and myocardium (myo). **p** magnified view of the EC (black square in **n)**. **r** In E12.5 OFT, the *Gpc6* signal (blue) in seen in the wall of the developing great vessels.

Next, we investigated the localization of the *Gpc6* mRNA in mouse embryonic hearts using *in situ* hybridization, utilizing a Gpc6-specific RNA probe (see Supplemental Figure 3 for specificity confirmation). In E10.5 and 11.0 hearts, the Gpc6 signal was detectable in the aortic bulb (arrow heads) but not in the heart itself (Figure 2c, E10.5 is shown). At E11.5, the Gpc6 signal was present in the outflow tract (OFT) and atrioventricular canal (AVC) of the heart (Figure 2d, f). Notably, a similar signal was observed for the *Pdgfra* gene, reported to be expressed in the mesenchymal cells occupying the endocardial cushions (31) (EC, Figure 2e). A day later, the expression of *Gpc6* was either greatly reduced (E12.5) or undetectable (E12.5+) in the AVC and proximal OFT cushions while still clearly detectable in the distal OFT (Figure 2g, i, j) where the great arteries (GA) develop. At the same time (E12.5), *Pdgfra* expression was still present in the AVC and proximal OFT but absent in developing GA (Figure 2h). The *Gpc6* expression in the walls of GA persisted until E14.5 and faded away after E15.5 (Figure 2k-m). Transverse histological sections of the Gpc6-stained hearts (E11.5 and 12.5, n=3 of each age) showed that at E11.5, *Gpc6* expression is clearly detectable in the EC mesenchyme but not in the endocardium or myocardium, and overlaps *Pdgfra* expression (Figure 2n-o). In E12.5 sections, the signal was detectable in the smooth muscle layer of the great vessels but not in the endothelium or adventitial layer (Figure 2r). We tested whether *Gpc6* was expressed during heart specification or in the cardiac neural crest cells (cNCC). No *Gpc6* signal was detected in the cardiac crescent (marked by *Isl1* expression in the second heart field, SHF (32)) at E7.5 and in the looping heart and SHF at E8.5 (Supplemental Figure 4). In E9.5 and E10 embryos, no signal was observed in the heart (h) or cNCC (arrow; *Crabp1* (33) in Supplemental Figure 4). These results strongly argue for the function of the Gpc6 gene being exclusive to the endocardial cushions.

While GPC6 was considered as the principal candidate gene, we also analysed the expression of Gpc5 (Supplemental Figure 5). Although *Gpc5* RNA was detected in mouse embryonic hearts, at E11.5 onwards, its widespread expression throughout the heart did not suggest specific function during OFT septation.

### Generation of the Gpc6 global KO mice

The Gpc6^tm2a (EUCOMM)Wtsi^ allele (subsequently referred to as Gpc6^tm2a^) was designed as “Knockout First, with Conditional Potential” (34), where the mRNA synthesis is terminated after exon 2 by a pA stop sequence (see allele schematics in Supplemental Figure 6a). However, Gpc6^tm2a/tm2a^ mice were viable, contradicting the previously described lethal phenotype in an alternative knockout strain (Gpc6^tm1Lex^), and did not exhibit major abnormalities. Further investigation confirmed the presence of the full length *Gpc6* mRNA in Gpc6^tm2a/tm2a^ tissues (Supplemental Figure 6c) indicating that the tm2a allele is a hypomorph. Crossing to a Keratin-14-Cre line was required to generate the “true” global *Gpc6* KO mice, in which we confirmed the expression of *Gpc6* to be absent (see Methods and Supplemental Figure 7b-e). Thereafter, we utilised the hypomorphic tm2a allele in compound heterozygous crosses with the newly generated KO allele to enable dose-response studies (see below).

### Phenotypic assessment of the Gpc6 global KO mice

Mice heterozygous for the KO allele (Gpc6^Δ/+^) were viable, healthy and fertile. Genotyping of the embryos obtained from a HET X HET intercross revealed the presence of all three expected genotypes with no statistically significant difference between observed and expected numbers, while the homozygous (Gpc6^Δ/Δ^) genotype was never found in adult mice (Supplemental Table 2).

In the newborn litters from heterozygous crosses, dead pups were found on the day of birth and confirmed to be the Gpc6^Δ/Δ^ genotype (Supplemental Figure 7b). They had a morphologically abnormal appearance, with pale skin, small body size, short snout (Supplemental Figure 8a, d), little or no air in the lungs (Supplemental Figure 8b, e) and cleft palate (Supplemental Figure 8c, f). Both fore- and hindlimbs were noticeably shorter in the KOs (Supplemental Figure 8g, h). An air bubble was found in the trachea (not shown), suggesting that the pups were alive at birth but died shortly after due to breathing difficulties.

Most importantly, the Gpc6 KO pups exhibited cardiac morphology defects closely related to human TOF, which were found with 100% penetrance. In the normal heart, the aortic inflow originates in the left ventricle and the aorta (Ao) then spirals around the PA. In Gpc6 KO hearts, both great arteries originated from the heart parallel to each other and the aortic root was always mal-positioned relative to the PA (Figure 3a, b). The conal myocardium, from which both arteries originated, appeared enlarged. Both Ao and PA originated from the right ventricle (Figure 3c, f), resulting in a double outlet right ventricle (DORV) phenotype. Dorsally, a perimembranous VSD was found (Figure 3d, e). The mitral and tricuspid valves were not affected (Figure 3e, h). The lumen diameter of the main pulmonary artery was significantly reduced compared to controls, as was the aortic lumen diameter (Figure 4a-c). No aorticopulmonary septal defect was observed (Figure 4d-e). Finally, the KO hearts exhibited RV myocardial hypertrophy (Figure 4f-h).

**Figure 3.**
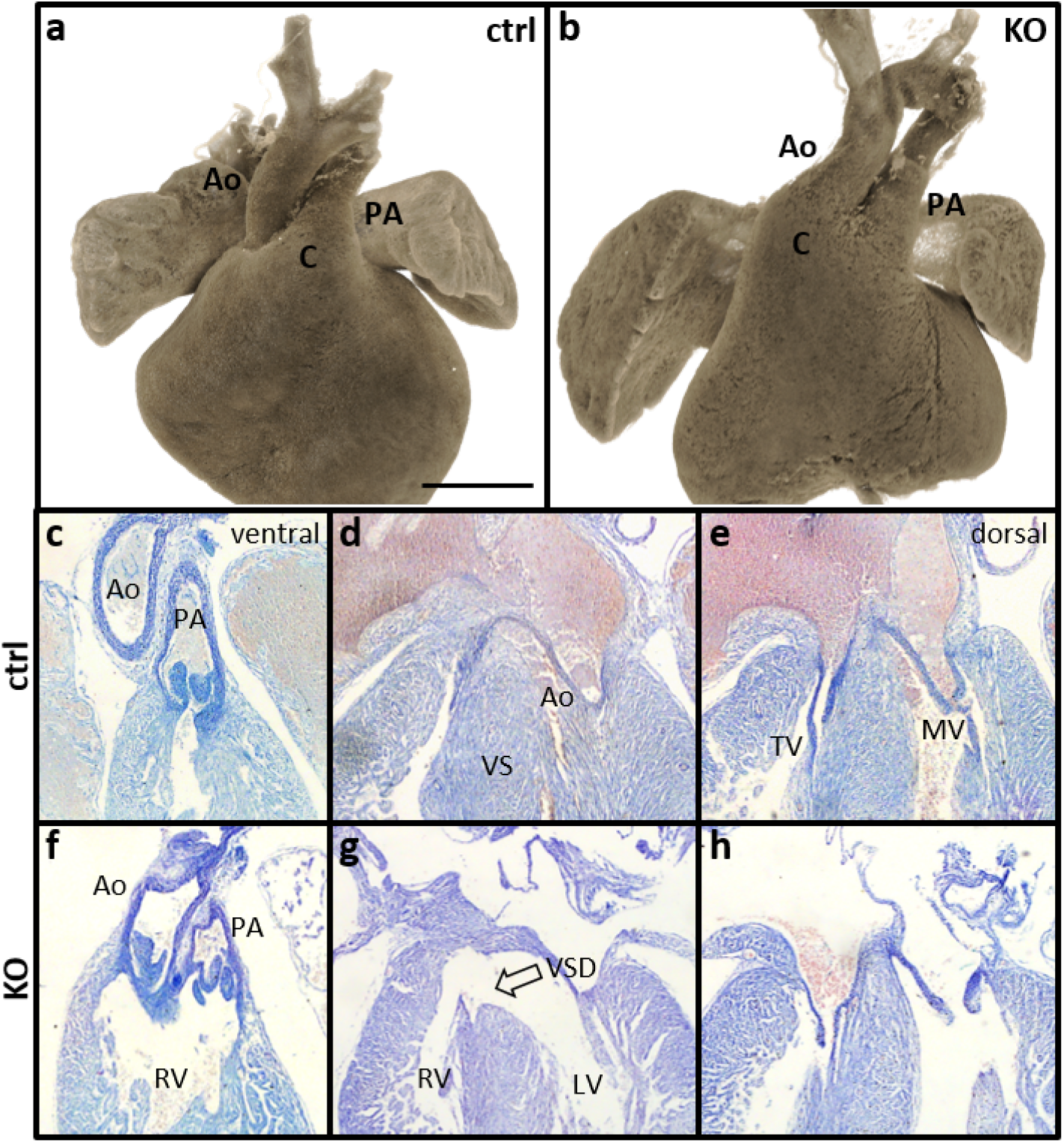
Phenotypic analysis of the Gpc6 KO hearts. **a-b** 3D reconstruction of the microCT scans of the control (**a**) and global KO (**b**) neonatal mouse hearts. Aorta is clearly identified by the presence of the aortic arch. Ao aorta, PA pulmonary artery, C conal myocardium. Scale bar 500 μm. Aorta in KO heartsis malpositioned relative to PA, and conal myocardium is distended. Aorta is clearly identified by the presence of the aortic arch. **c-h** Histological examination of the neonatal hearts confirmed that both great arteries originate from the right ventricle (RV; **c** vs **f**) and dorsally, ventricular septal defect (VSD) was found (**d** vs **g**). No serious anomalies were observed in mitral (MV) and tricuspid (TV) valves (**e** vs **h**). LV left ventricle.

**Figure 4.**
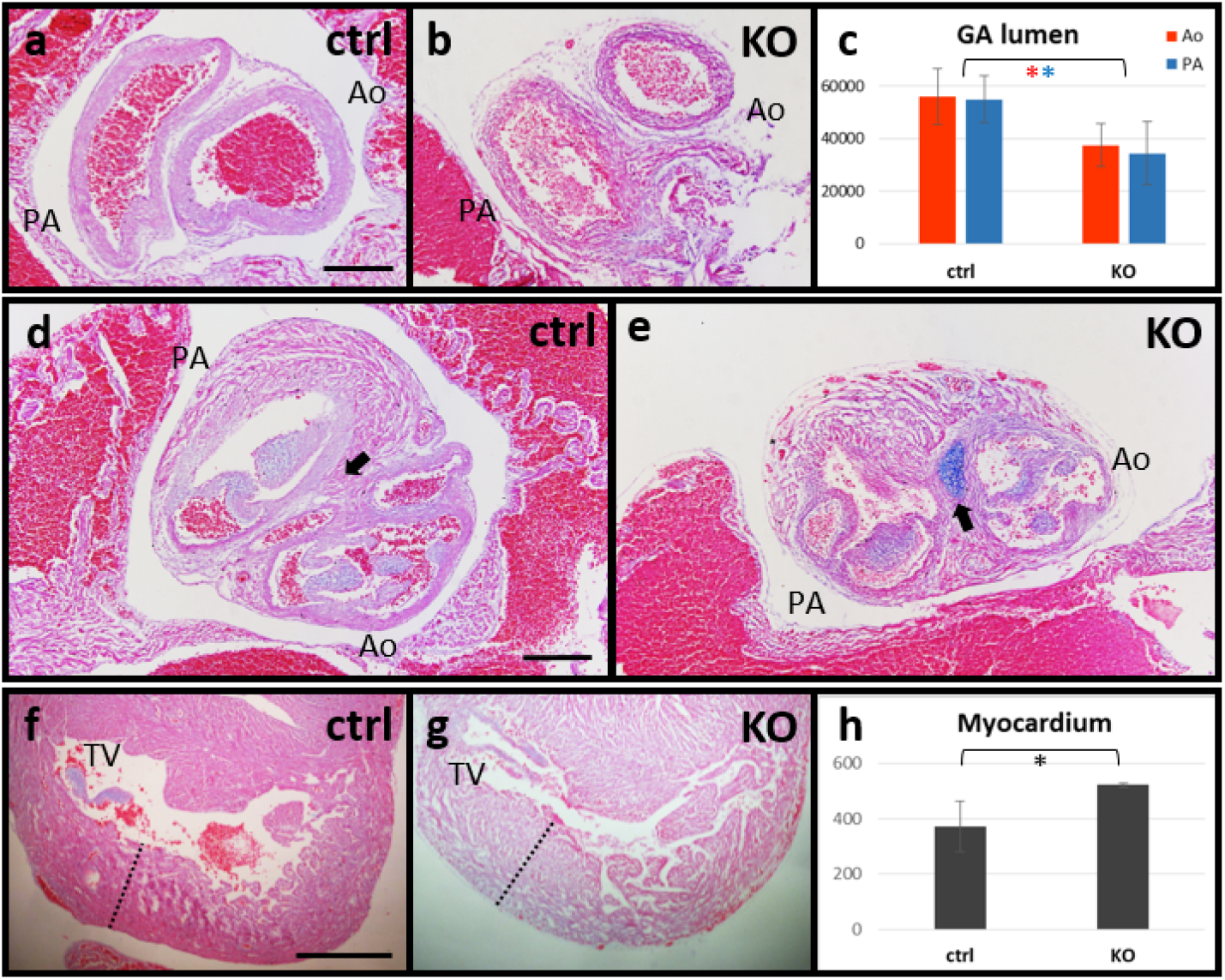
Arterial and right ventricular phenotypes in the Gpc6 KO hearts. **a-c** Gpc6 KO neonatal hearts exhibited reduction in both the pulmonary and aortic lumen areas (P = 0.0235 for Ao and 0.0148 for PA). **d-e** No aortico-pulmonary septal defect was observed in the KO hearts (black arrow points to intact AP septum). **f-h** KO hearts exhibited right ventricular hypertrophy indicated by thickening of the myocardial wall (dashed line, P = 0.0156). Asterisks in c and h highlight the statistically significant differences (Student’s t-test). TV tricuspid valve. Scale bar: 200 μm for **a-b** and **d-e**, 500 μm for **f-g**.

Of note, although the *lacZ* transgene was not removed from the *Gpc6^tm2a^* allele prior to generation of the KO mice, DORV and other developmental defects were only observed in homozygous KO animals (2 x *lacZ* copies). No such abnormalities were found in *Gpc6^tm2a/tm2a^* mice (2 x *lacZ* copies) as well as in either *Gpc6^tm2a/+^* or *Gpc6^Δ/+^* animals (1 x *lacZ* copy), suggesting that the *lacZ* transgene has no noticeable influence on the phenotype.

### Embryological origin of cardiac defects in Gpc6 KO mice

We next identified when the Gpc6 KO cardiac defects develop. In E11.5 embryonic hearts, no phenotypic differences were observed between WT and HOM samples (Figure 5a, e). At E12.5 and E13.5, the OFT morphology appears abnormal in KO hearts (Figure 5b vs f, c vs g): both developing Ao and PA are connected to the right ventricle and vessel lumens run parallel to each other instead of the normal spiralling appearance. By E15.5 this becomes more apparent and resembles the neonatal phenotype (Figure 5d, h).

**Figure 5.**
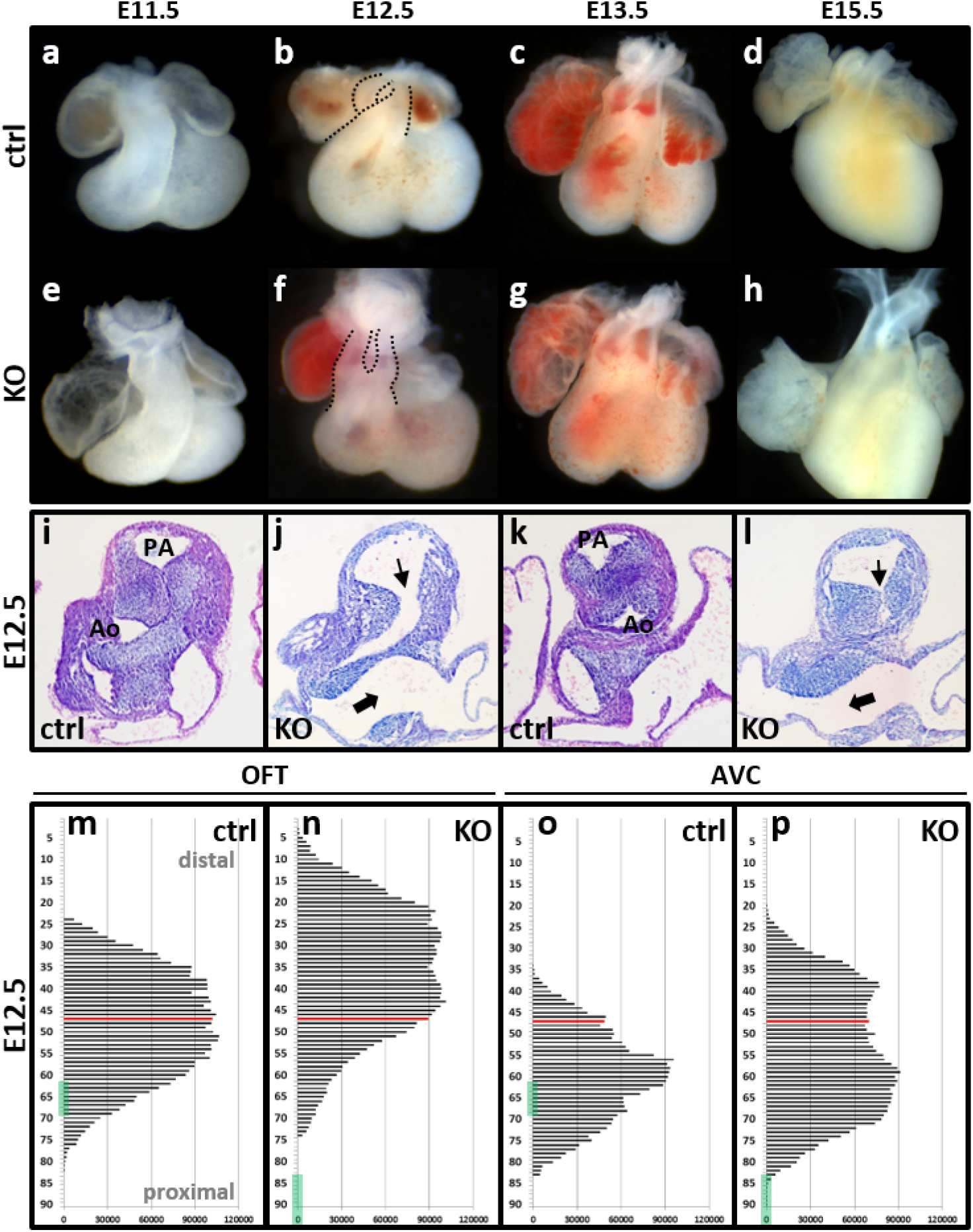
Gpc6 KO produces abnormal arterial alignment and endocardial cushion perturbation. Consistent with Gpc6 gene expression time course, no visible abnormalities were observed in E11.5 heart (**a** vs **e**). By E12.5, delayed and abnormal rotation of the common OFT is detectable in KO hearts compared to controls (**b** vs **f**, great arteries are outlined). At E13.5 (**c** vs **g**) and 15.5 (**d** vs **h**), the DORV phenotype becomes more pronounced, with great arteries growing parallel to each other. **i-l** Histological examination of E12.5 hearts. Transverse sections of the proximal (**i** vs **j**) and distal (**k** vs **l**) OFT show smaller EC area and delayed formation of the OFT (thin arrow) and AVC (thick arrow) septa. For further analysis, the area occupied by cushions in each section was measured (n=3 of each genotype) and the distribution of the EC volume within the heart showed notable differences between control and KO samples (**m-p**). The Y axis represents sections in distal to proximal direction with bars showing the size of the area occupied by EC in each section (in square pixels). In both the OFT and AVC, EC were shifted distally relative to the morphological reference level (selected as the first section where OFT is fully separated from the ventricular myocardium, red line); the KO cushions also started above the upper edge of the ventricular septum (green bar), while in control samples they start deeper in the ventricle (OFT, **m** vs **n**; AVC, **o** vs **p**).

Next, we assessed the morphology of the endocardial cushions at E12.5. There was severely delayed septation of both the AV canal and OFT (Figure 5i-l). There was no statistically significant difference in the volume of the EC between KO and WT samples (Figure 5m-p): in the OFT, 3.55×10^6^ in WT vs 3.98×10^6^ square pixels in KO, P=0.42; in the AVC, 2.28 x10^6^ in WT vs 3.51 x10^6^ in KO, P=0.107 (Chi-Square test). However, a clear difference in the cushion morphology was noted. Firstly, in both the OFT and AVC, cushions appeared elongated. Secondly, in the OFT, the KO cushions were shifted in the proximal to distal direction and the distance between the reference section and the ventricular septum edge was increased. Thus, the EC in controls originate deeper in the ventricle, under the edge of the VS, while it starts above the VS in the KO. In the AVC, the KO cushions were shifted deeper into the ventricles but still originated above the VS edge while in the WT hearts, the VS edge lies in the middle of the cushions.

### Gpc6 knock-down exhibits dose-dependent effect on the phenotype

We hypothesised that further reduction of the Gpc6 transcript abundance below 50% (in Gpc6^Δ/+^) might cause a milder and/or less penetrant phenotype than the DORV observed in Gpc6^Δ/Δ^. Gpc6^Δ/tm2a^ animals retain about 20% of the Gpc6 transcript (Supplemental Figure 7e). Newborn pups were alive, with normal appearance and no cleft palate (n=13). Five of 10 hearts investigated had noticeably thinner perimembranous VS (Figure 6a vs b-c) and two exhibited complete sub-aortic perimembranous VSD (Figure 6d). One heart displayed a TOF-like DORV, with overriding aorta and sub-aortic perimembranous VSD (cKO-4 in Figure 6e-i); aortic lumen size appeared within the normal range (Figure 4c, 54787 μm^2^), while PA lumen was reduced (43713 μm^2^).

**Figure 6.**
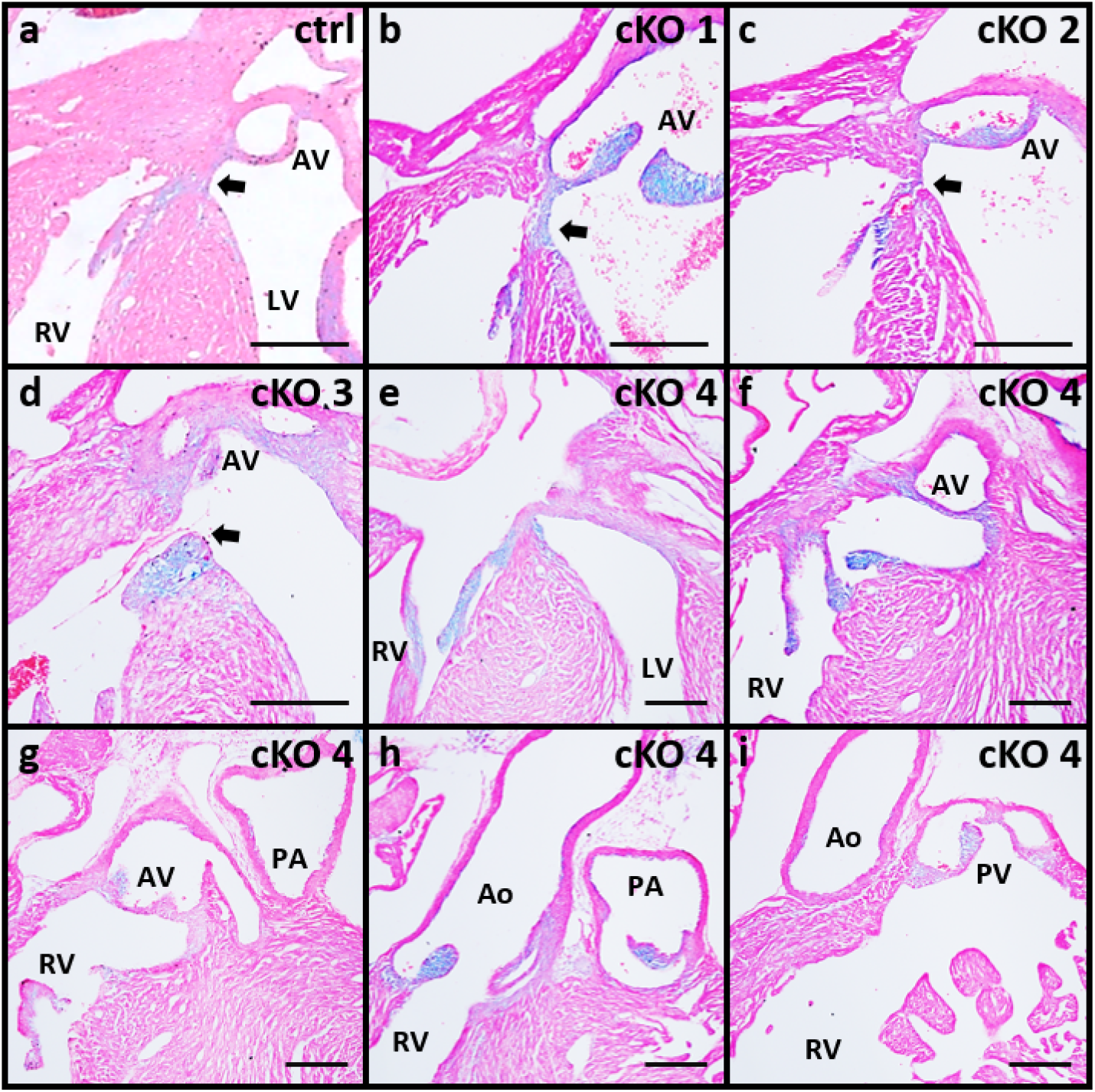
Phenotype of compound Gpc6 Δ/tm2a neonatal hearts. Histological examination of the neonatal hearts of compound Gpc6^Δ/tm2a^ genotype (cKO). Arrow points to perimembranous ventricular septum (mVS). **a** Intact mVS in control sample. **b-c** Thin mVS in 2 examples of cKO hearts. **d** mVS failure to close in cKO-3. **e-i** Series of sections of cKO-4 heart (dorsal to ventral). Aorta (Ao) is connected to both left (**e**) and right (**f-h**) ventricles, while pulmonary artery (PA) originates from right ventricle (RV). LV left ventricle, AV aortic valve, PV pulmonary valve. Scale bar 200 μm.

### Tissue-specific Gpc6 KO recapitulates TOF phenotype and implicates EC mesenchymal cell defect

The EC mesenchyme consists of two cell populations, namely endoMES cells (derived from the endocardium via the endocardial-to-mesenchymal transition, EndoMT), and cardiac NCC (35). To test whether Gpc6 function is required in either cell type or both, we generated the tissue-specific gene KOs utilizing Tie2-Cre (for endoMES) and Wnt1-Cre (for NCC). EndoMES KO animals were viable with no detectable detects. NCC KO mice died neonatally with airless lungs, and exhibited cleft palate but no cardiac anomalies. In contrast, homozygosity for the combination of the two conditionally deleted genotypes resulted in a sub-aortic VSD accompanied by overriding aorta (Supplemental Table 3 and 4, Figure 7), a phenotype on the same anatomical spectrum as the TOF-type DORV observed in the wholebody KO. This phenotype is less extreme regarding the arterial malposition and indeed closer to the typical human TOF phenotype in this respect than was the whole-body KO. No change in the size of GA or myocardium was observed, consistent with an overall milder phenotype. These results together with the cushion abnormalities observed, indicate that the *Gpc6* KO cardiac phenotype results from a defect in the function of the endocardial cushion mesenchyme.

**Figure 7.**
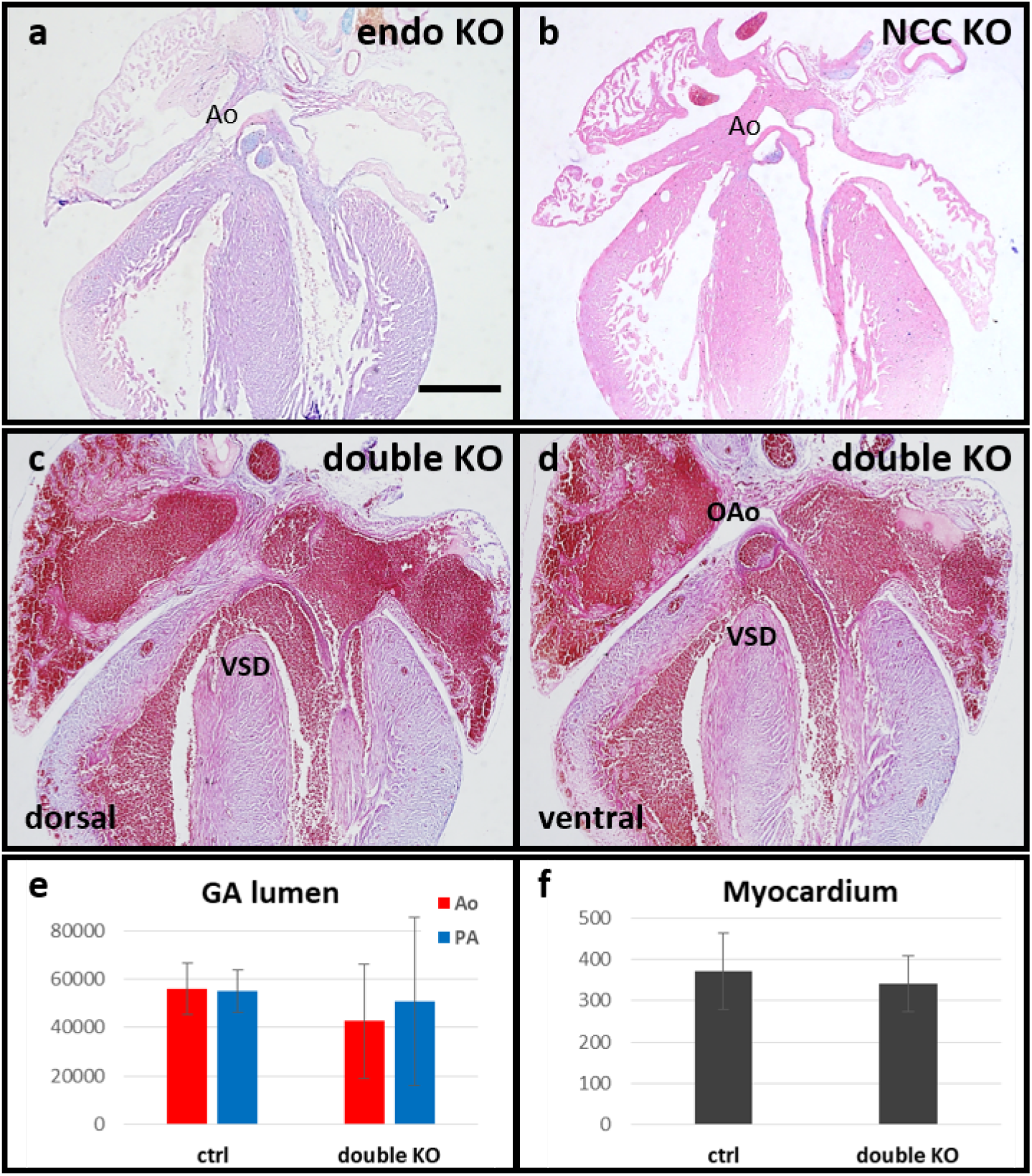
Cardiac defects in animals with cell-type specific deletion of the Gpc6 gene. Coronal sections of the neonatal mouse hearts. **a** Endocardial lineage-specific deletion produced with Tie2-Cre (will affect all EC mesenchymal cells derived from the endocardium via EMT). **b** Neural crest cell lineage-specific deletion produced with Wnt1-Cre. **c-d** Combined deletion of Gpc6 in both (endo+NCC) mesenchymal cell populations results in a sub-aortic VSD accompanied by the overriding aorta (OAo). No significant changes were observed in double KO hearts for the lumen crosssection size of either aorta or pulmonary artery (Ao and PA in **e**, correspondingly), as well as in right ventricular myocardium thickness (**f**) (Student’s t-test). No cardiac defects were found in [Tie2-Cre, Wnt1-Cre] controls. Values in μm^2^ in **e** and μm in **f**. Scale bar 500 μm.

### Gene expression analysis

After establishing the functional importance of Gpc6 in EC morphogenesis, we analysed the changes in expression of key genes involved in this process in E11.5+ outflow tracts (WT controls and KO samples, in 3 biological repeats). First, we investigated the rate of proliferation and apoptosis. Proliferation markers (Ki67, Tpx2, Top2A) and pro- (Casp2 and Casp8) and anti-apoptotic (Bcl2) genes (Table 1), were not different in wildtype and KO outflow tracts. These findings are consistent with our observation that there was no change in the EC volume. Next, we looked at the expression of key genes involved in EndoMT, and EC cells differentiation and migration. Out of 8 genes investigated, 2 genes involved in EMT (Snai1, Adam19) and 4 genes involved in cell differentiation and migration (Adam19, Pdgfrb, Rxra, Tbx20) were significantly upregulated. This, together with our previous finding of severely mis-shaped EC, suggests that abnormal mesenchymal cell migration is likely to play a significant role in the observed phenotype. Finally, we analysed non-canonical Wnt/PCP signalling based on the overrepresentation of the genes of this pathway in causative genes for both DORV and TGA phenotypes in the mouse (Supplemental Figure 9) and their role in directional cell migration during organ formation. Of 7 components tested, two (signal transducer Dvl1 and trans-membrane receptor Ptk7) were significantly upregulated (Table 1).

**Table 1.** Differential gene expression analysis.

| Gene | Gene function | Cardiac phenotype | Fold Change | SD | P-value |
| --- | --- | --- | --- | --- | --- |
| *Proliferation* |  |  |  |  |  |
| *Ki67* | protein present in G<sub>1</sub>, S, G<sub>2</sub>, and M but absent in G<sub>0</sub> |  | 1.089 | 0.119 | 0.2402 |
| *Tpx2* | spindle assembly factor during mitosis |  | 1.074 | 0.038 | 0.0511 |
| *Top2A* | a core component of mitotic chromosomes and important for establishing mitotic chromosome condensation |  | 1.053 | 0.117 | 0.4654 |
| *Apoptotic* |  |  |  |  |  |
| *Bcl2* | anti-apoptotic factor |  | 0.857 | 0.192 | 0.3517 |
| *Casp2* | mediate cellular apoptosis through the proteolytic cleavage of specific protein substrates |  | 0.998 | 0.056 | 0.9576 |
| *Casp8* |  |  | 1.070 | 0.089 | 0.3298 |
| *Endocardial Cushions genes* |  |  |  |  |  |
| *Adam19* | proteolytic regulation of ErbB ligands critical for EMT; regulate differentiation of the cNCC in the OFT | DORV, OAo, VSD, stenosis | **1.301** | **0.112** | **0.0191** |
| *Pdgfra* | regulates correct migration of the cNCC-derived fibroblasts within the OFT |  | 1.428 | 0.237 | 0.0629 |
| *Pdgfrb* | regulates MC migration and ECM production | DORV, OAo, VSD, AVSD, PTA | **1.303** | **0.182** | **0.0462** |
| *Rxra* | Retinoic acid receptor important for correct morphogenesis of EC around E12.5 | DORV, PTA, PS, AVSD | **1.155** | **0.055** | **0.0161** |
| *Snai1* | key transcriptional factors required for EMT initiation in the endocardium | abnormal AVC, hypoplastic AV cushions, reduced EMT | **1.505** | **0.215** | **0.0292** |
| *Snai2* |  |  | 1.301 | 0.307 | 0.2381 |
| *Twist1* | key transcriptional factor in EC mesenchyme; induces mes cells proliferation and promotes migration |  | 1.462 | 0.267 | 0.0705 |
| *Tbx20* | positively regulates EC mesenchyme proliferation and migration; downstream of Twist1 | hypoplastic AV cushions | **1.664** | **0.254** | **0.0010** |
| *Wnt-PCP pathway* |  |  |  |  |  |
| *Cnn2* | an effector of Wnt-PCP signalling in migrating cNCC |  | 0.961 | 0.022 | 0.2247 |
| *Daam1* | an effector of noncanonical Wnt-PCP signalling | DORV, VSD | 0.971 | 0.281 | 0.9262 |
| *Dvl1* | key component of the signal transduction in Wnt signalling |  | **1.436** | **0.033** | **0.0057** |
| *Ptk7* | co-receptor of the Wnt-PCP pathway | DORV, VSD, dilated PA | **1.256** | **0.041** | **0.0243** |
| *Vangl2* | transmembrane protein with regulatory role in Wnt-PCP pathway | DORV, VSD | 1.155 | 0.246 | 0.5918 |

## Discussion

Taken together, our results show that *Gpc6* is the gene overwhelmingly likely to be responsible for the GWAS association signal with TOF at the 13q31 locus. We demonstrated that the timing and location of *Gpc6* expression in the developing heart are consistent with a role in outflow tract septation; specifically, in the endocardial cushion mesenchyme. We established a clear dose-response relationship between *Gpc6* expression and OFT morphology in the mouse. Animals with complete absence of *Gpc6* had TOF-type DORV with 100% penetrance, while homozygotes for the hypomorphic tm2a allele expressed Gpc6 at around 50% of normal levels, and did not have cardiac defects. Compound heterozygote animals for the tm2a and knockout alleles expressed around 20% of normal *Gpc6* levels, and had incompletely penetrant TOF-like phenotypes with less severe malposition of the aorta relative to the pulmonary artery. Through tissue-specific knockout we confirmed our hypothesis, developed from our expression and anatomical data, that the endocardial cushions are the key developmental structures susceptible to reduced Gpc6 levels. Gene expression analyses indicated genes involved in EC biology and non-canonical Wnt signalling were dysregulated as a result of *Gpc6* knockout.

Several lines of evidence suggested that out of the two closest coding genes at chr13q31 locus (the only genes within ^~^2 Mb of the GWAS signal), GPC5 and GPC6, the latter was the more plausible candidate for TOF. First, based on population genomic data, GPC6 is less tolerant to mutations (missense, LOF and CNVs) than GPC5, indicating that the disruption of GPC6 function is more likely to cause developmental defects. Second, the rs7982677 haploblock is located in the same TAD as the whole of the GPC6 gene but not the GPC5 gene. Given that a TAD typically contains both genes and their *cis*-acting regulatory sequences (36), we reasoned that TOF-associated variants are more likely involved in the regulation of GPC6 but not GPC5 expression. Third, a number of chromosomal features link LD region to the GPC6 promoter; similar links were not found for the GPC5 promoter. These data argue for the existence of important regulatory elements within the GWAS locus, regulating GPC6 expression. None of the variants in the rs7982677 haploblock are identified as significant eQTLs in GTEx. However, this resource only encompasses adult tissues, and absence of an eQTL in these tissues does not rule out a developmentally specific influence of the SNP on expression – indeed such stage-specific eQTLs were recently identified in developing cardiomyocytes (37). The non-coding RNA GPC5-AS2, located in the LD region, was not detected in human embryonic heart at CS12-22 when TOF defects develop, which ruled it out as a TOF candidate.

In our study, global Gpc6 deletion resulted in TOF-type DORV, which is often considered to be the most severe type of TOF. Likewise, compound heterozygosity for the tm2a and deletion alleles, or tissue-specific Gpc6 deletion in the EC mesenchyme, produced the phenotype of a sub-aortic VSD with the aortic origin overriding RV and LV (that is, less severely malpositioned), which is closer to the typical human TOF phenotype. We explain the phenotypic difference between the global and mesenchymal KO by a likely incomplete efficiency of the tissue-specific Cre transgenes, which is previously recognised in such models (38). Thus, we showed either whole-body or cushion mesenchyme-specific Gpc6 deletion in mice leads to outflow tract defects clearly related to TOF, and established a dose-response relationship (Figure 8a). Our observations Support the hypothesis that a genetic defect in a single gene can cause a range of OFT malformations, which in clinical practice are considered as individual CHDs (39, 40).

**Figure 8.**
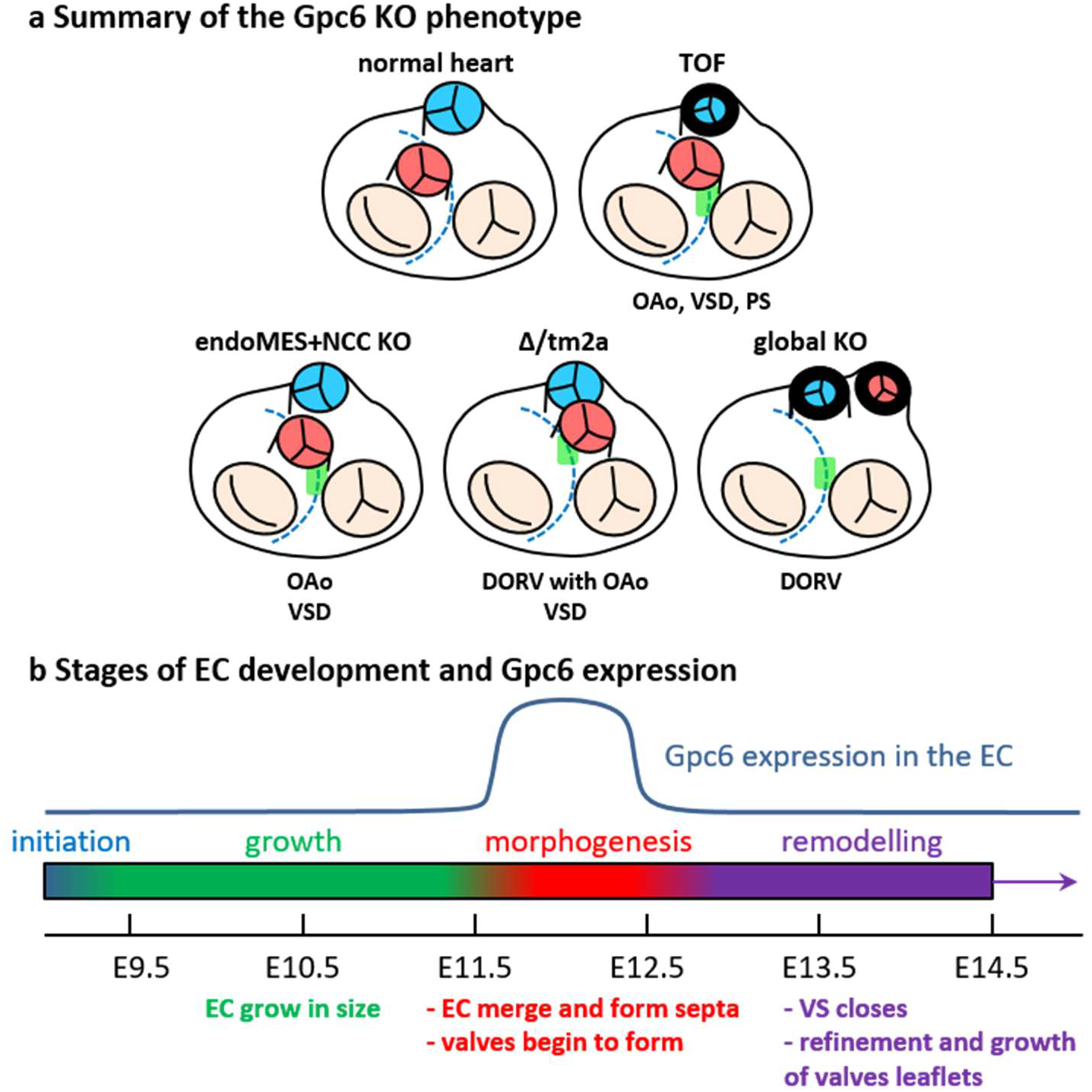
Gpc6 in cardiac septation. **a** Heart schematics (top view) summarise the cardiac defects data. Pulmonary artery in blue, aorta in red. Dashed line represents ventricular septum. Green bar denotes the ventricular septal defect. OAo overriding aorta, VSD ventricular septal defect, DORV double outlet right ventricle. **b** The time line of the cardiac septation in mouse embryo with four stages of endocardial cushions (EC) development: initiation, growth, morphogenesis and remodelling. Blue line represents the expression of the Gpc6 gene in the EC (excluding the aortic bulb).

In humans, homozygous LOF mutations in GPC6 have been implicated in the very rare condition Omodysplasia-1 (41) (OMOD1, OMIM:258315; only 27 cases reported), which is characterized by severe congenital micromelia and facial dysmorphism. Several OMOD1 patients have been described with CHD phenotypes, including truncus arteriosus, VSD, and pulmonary stenosis (4/27 cases described to date) (42, 43). This evidence further supports the involvement of GPC6 in human CHD as a susceptibility factor which interacts with other genetic and environmental factors to cause sporadic, non-syndromic TOF. Of note, patients in the original TOF GWAS identifying the 13q31 region were non-syndromic and without extracardiac malformations. The mechanisms whereby some classes of variation in *GPC6* cause limb defects and occasional cardiac defects in patients, and others cause congenital heart disease in the absence of limb malformations, remain to be explored.

Glypicans are heparan sulfate proteoglycans that are bound to the outer surface of the plasma membrane by a glycosyl-phosphatidylinositol anchor. They are evolutionarily conserved proteins; there are six family members in vertebrates (44). Glypicans are highly expressed in a stage- and tissue-specific manner during embryogenesis, suggesting an involvement in morphogenesis (45). They act as co-receptors whose main function is to regulate key developmental pathways including Wnt, Shh, FGF and BMP signalling (44), all of which have been implicated in cardiac septation and whose disruption may lead to CHDs (46). So far, two glypicans have been linked to CHDs. *Gpc3* mutations have been identified in Simpson-Golabi-Behmel syndrome (47), and *Gpc3*-deficient mice display a high incidence of congenital heart defects including VSD, common atrioventricular canal and DORV (48). In zebrafish, gpc4 deletion causes a reduction in size of both atrium and ventricle (49). Our results argue for a main function of *Gpc6* in EC morphogenesis rather than specification or growth (Figure 8b). Moreover, the data suggest that Gpc6 regulates cell migration during cushion morphogenesis, but not mesenchymal cells proliferation or apoptosis, as no change in corresponding marker genes was observed; consistent with this, the EC have normal size but are abnormally shaped and distributed in KO hearts. The key process for correct EC morphogenesis is coordinated cell movement. The non-canonical Wnt, planar cell polarity (Wnt-PCP) signalling pathway has been heavily implicated in this process. Mutations in different components of the pathway lead to a range of CHDs, including DORV, TGA and PTA (50). Consistently, we found key members of Wnt-PCP pathway to be upregulated in the Gpc6 KO hearts. It has long been known that glypicans modulate Wnt signalling, acting as either positive or negative regulators (51, 52). The molecular mechanism of such modulation has recently been described (53). As such, we propose that Gpc6 plays a crucial physiological role in the EC morphogenesis by attenuating local Wnt-PCP signalling. Thus, loss of Gpc6 may lead to overactivation of the pathway and abnormal mesenchymal cell migration. Further research will be required to confirm this.

Taken together, our results implicate GPC6 as the causal gene for Tetralogy of Fallot in the 13q31 region. No non-syndromic condition has to our knowledge been associated with Glypican genes thus far, and this is the first identification of a causative CHD gene from a GWA study. Though we have identified the endocardial cushions as the key structures in which GPC6 contributes to heart development, further work will be required to delineate the signalling events whereby GPC6 affects their morphology. Thus far only a relatively small number of CHD patients have been studied by GWAS; these results illustrate the feasibility of discovering causative genes for CHD through GWAS. With current technology GWAS could be carried out by whole genome sequencing, providing access to both rare and common variants. A more complete knowledge of CHD genetics could have downstream applications in genetic counselling for families affected with CHD.

## Methods

### Mice

The Gpc6^tm2a (EUCOMM)Wtsi^ (Gpc6 with floxed exon 3) mouse strain was generated by the Wellcome Trust Sanger Institute (MGI:4453713) and obtained from the Infrafrontier repository (www.infrafrontier.eu). The Gpc6 KO mice were produced with Keratin14-Cre females (MGI:2652655). Tie2-Cre (B6.Cg-Tg (Tek-cre)1Ywa/J) mice were used for endocardial *Gpc6* deletion. Gpc6 deletion in neural crest cells was achieved with Wnt1-Cre mice (54). Embryos and new-born pups were genotyped by PCR of genomic DNA (see Supplemental Table 1 for primer sequences and products sizes). To confirm their identity, PCR products were extracted from the agarose gel with ISOLATE II PCR and Gel Kit (Bioline) and Sanger sequenced.

### Expression analysis

To analyse gene expression, mouse hearts (embryonic days E9.5-15.5) were dissected in room temperature PBS. Human embryonic heart cDNA samples of different Carnegie stages (CS) were obtained from the MRC/Wellcome Human Developmental Biology Resource (hdbr.org). Mouse tissues were stored at −20°C in RNAlater solution (Ambion) for RNA extraction or fixed in 4% paraformaldehyde in PBS and stored at 4°C for in situ hybridization. RNA was isolated from tissue samples using Isolate RNA mini kit (Bioline) and cDNA was prepared with Tetro cDNA Synthesis kit (Bioline). qPCR was performed on Applied Biosystems ViiA™ 7 Real-Time PCR System with Sybr Green, and fold change was calculated using the ΔΔCt method.

### In situ hybridization

Whole mount *in situ* hybridization was carried out as previously described(55). The DNA templates for *Gpc6, Pdgfra* and *Isl1* probes were generated by PCR amplification of the corresponding gene’s coding sequence fragments from the mouse heart cDNA. DIG-labelled RNA probes were produced by *in vitro* transcription with T3-RNA polymerase (Promega). The *Crabp1* probe template was amplified from a plasmid containing *Crabp1* cDNA sequence. Images were acquired on a Leica S6D microscope with Leica MC170HD camera.

### Histology

Tissues were fixed in 4% paraformaldehyde in PBS for at least overnight and dehydrated through the 25-50-75% ethanol series. For histological examination, samples were then embedded in paraffin (Paraplast, Sigma), sectioned at 7 μm thickness, stained with 1% eosin Y and Alcian Blue. Photographs were taken on Olympus BX53 microscope with an Olympus DP74 camera. All measurements were performed using ImageJ software (56).

### micro-CT scanning

Neonatal hearts were fixed, mounted in paraffin and scanned on the ZEISS Xradia 520 Versa 3D X-ray microscope with following conditions: source energy 60kV, 5W; 4x objective; 3.6 microns per pixel size and a field of view of 3600 x 3600 microns, with 2s exposure per projection. 3D models were rendered with Drishti (57).

### Statistics

The viability of the KO embryos and adult mice was assessed by comparing numbers of the observed and expected genotypes. The statistical significance of the deviation from the expected Mendelian ratio was calculated with Chi-Square test (two-tailed P value; df 2 for global KO, 3 for tissue-specific KO). Gene expression in E11.5+ outflow tracts was quantified with qPCR (WT controls and KO samples, 3 biological repeats each) and the statistical significance was calculated with unpaired Student’s t-test (df = 3).

### Study approval

Mice were maintained in the Biological Service Facility (University of Manchester, UK) with local ethical approval and according to UK Home Office requirements (project P3A97F3D1).

## Author contributions

GT and BK conceived the project. GT designed research studies, conducted experiments, acquired data, analysed data, prepared figures, and wrote the original draft of the manuscript. AC assisted with the design of the qPCR, conducted qPCR experiments, and analysed data. KEH assisted with the design of the mouse experiments. KEH and BK acquired funding, supervised the project, and reviewed and edited the manuscript.

## Acknowledgements

Keratin14-Cre females were kindly provided by Prof Adrian Woolf, The University of Manchester (UoM). Tie2-Cre mice were kindly donated by Dr Delvac Oceandy, UoM. Wnt1-Cre mice were kindly provided by Prof Giulio Cossu, UoM. The plasmid containing *Crabp1* cDNA sequence was kindly provided by Prof Nicoletta Bobola, UoM. This study makes use of data generated by the DECIPHER community. A full list of centres who contributed to the generation of the data is available from https://deciphergenomics.org/about/stats and via email from. Funding for the DECIPHER project was provided by Wellcome. Those who carried out the original analysis and collection of the data bear no responsibility for the further analysis or interpretation of the data. We acknowledge the Engineering and Physical Science Research Council (EPSRC) for funding the Henry Moseley X-ray Imaging Facility which has been made available through the Royce Institute for Advanced Materials through grants (EP/F007906/1, EP/F001452/1,EP/I02249X, EP/M010619/1, EP/F028431/1, EP/M022498/1 and EP/R00661X/1), and personally Julia Behnsen (UoM) for performing CT scanning of the mouse hearts. The human embryonic and foetal material was provided by the Joint MRC/Wellcome Trust (grant# MR/R006237/1) Human Developmental Biology Resource (http://hdbr.org). We thank Dr E J Cartwright for support of the animal work. This work was supported by British Heart Foundation (UK) Programme Grant RG/15/12/31616 (to BK and KEH) and BHF Personal Chair CH/13/2/30154 (to BK).

## Notes

Conflict of interest: The authors have declared that no conflict of interest exists.

### Competing Interest Statement

The authors have declared no competing interest.

