## Supplemental data for "Glypican-6 deficiency causes dose-dependent conotruncal congenital heart malformations through abnormal remodelling of the endocardial cushions"

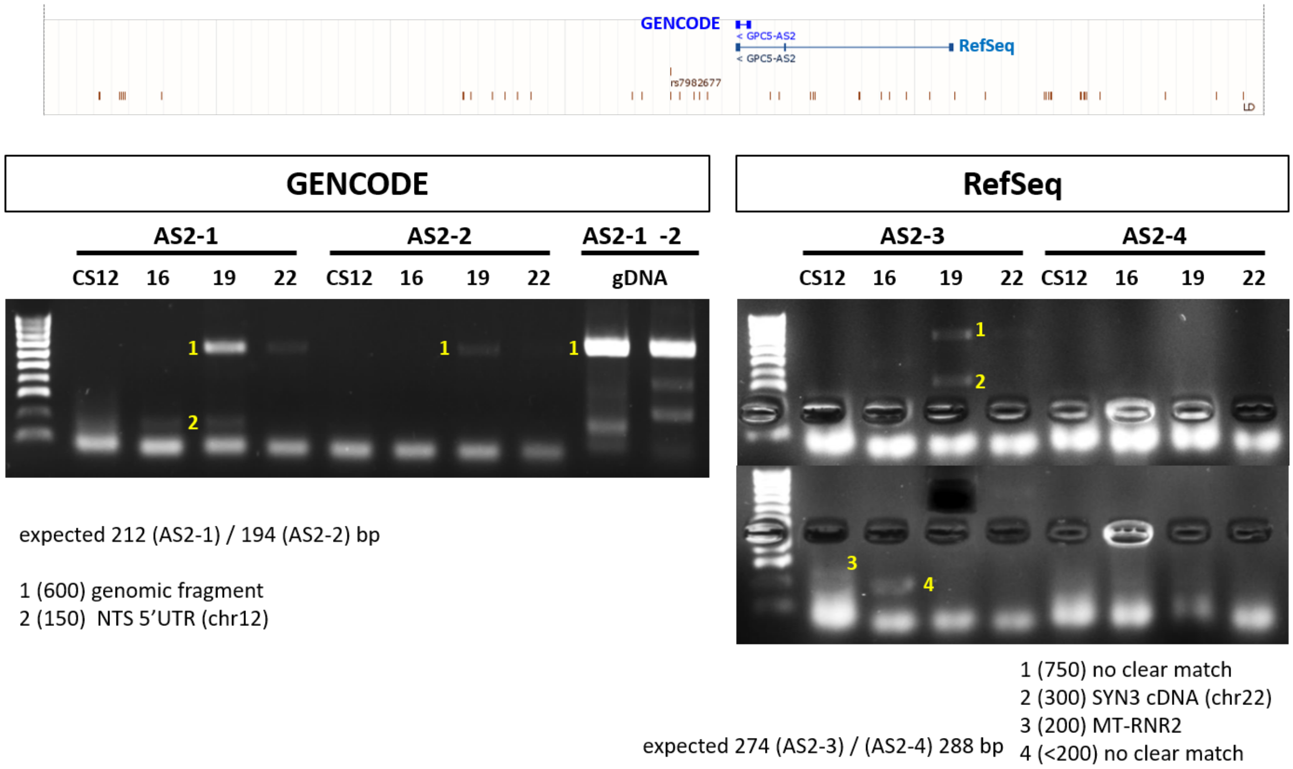


**Supplemental Figure 1. Analysis of the Gpc5-AS2 RNA expression in human heart.**

**a** The overview of the chromosomal region containing TOF associated haploblock (individual SNPs are shown as brown bars). Two annotations of the GPC5-AS2 non-coding RNA are shown (GENECODE and RefSec). **b** The analysis of GPC5-AS2 expression in human embryonic heart tissue. CS - Carnegie stages of human embryonic development. gDNA - control PCR with genomic DNA. AS2-1 (-2, -3, -4) four sets of primers used in expression analysis (see Sup Table 1). The sequencing results of the products 1-6 are shown under corresponding gels.

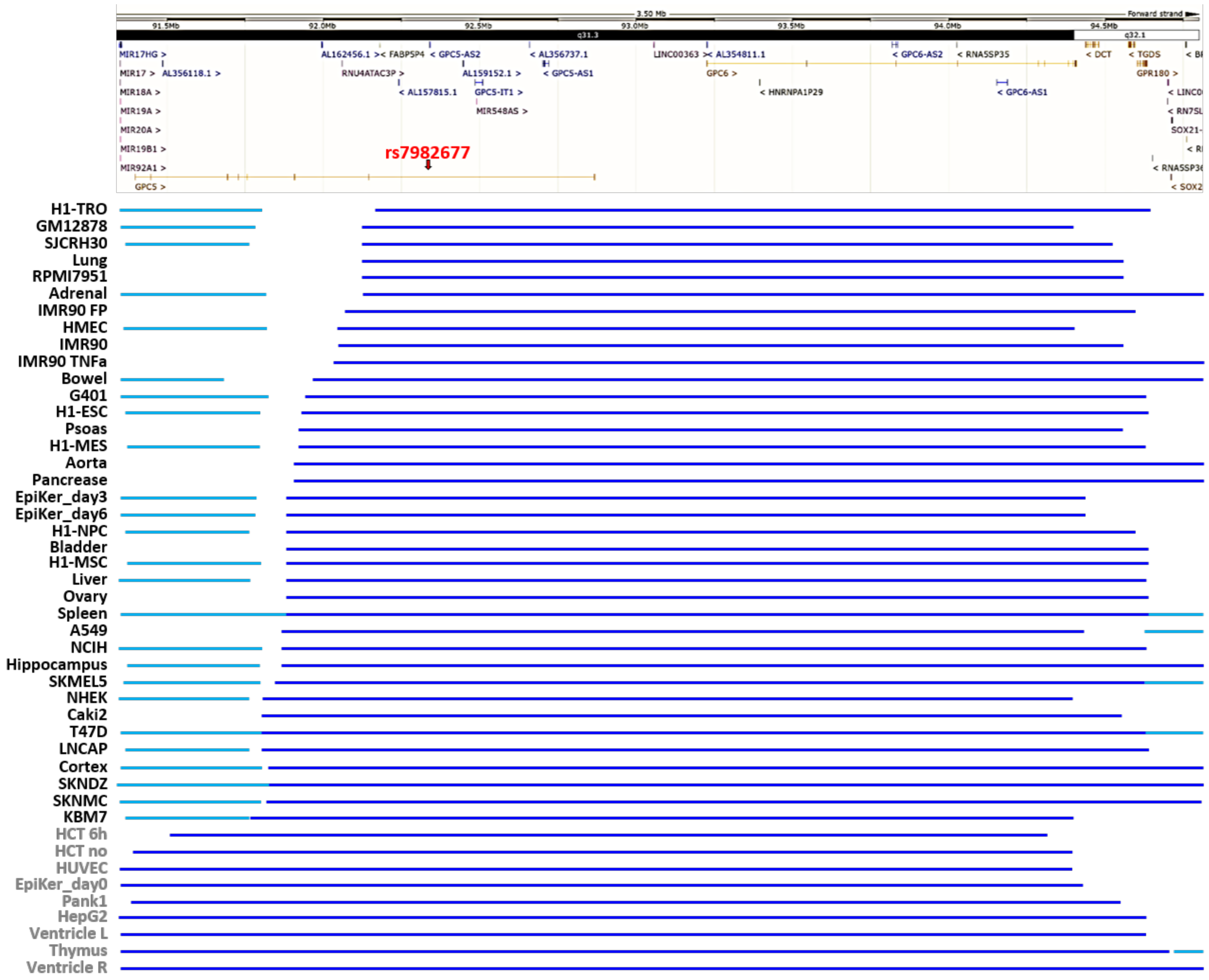


**Supplemental Figure 2. Structure of topologically associated domains (TADs) in chr13q31 locus.**

Top: the overview of the 3.5 Mb chromosomal region containing both GPC5 and GPC6 genes. The location of the rs7982677 variant is shown with red arrow.

Bottom: TADs in 46 cell lines/tissues. Grey labels - cell lines/tissues where single domain encompasses both GPC5 and GPC6 genes. Black labels indicate that the domain only includes the whole of the GPC6 gene. Alternating domains are shown with light/dark blue lines.

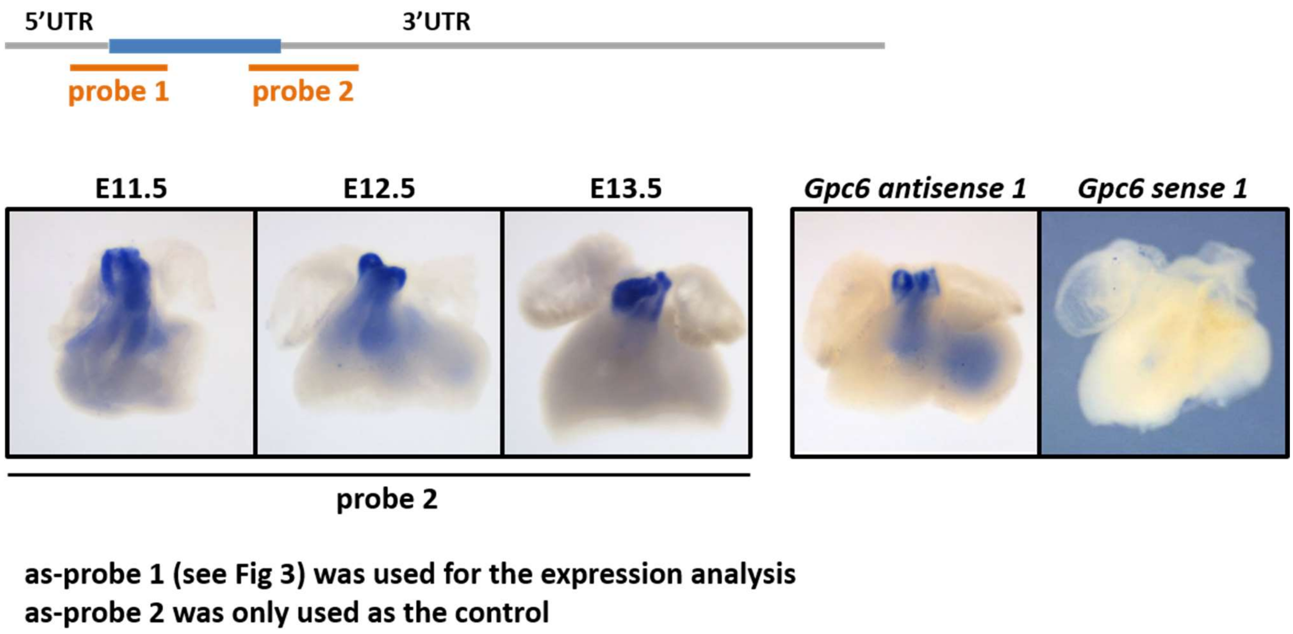


**Supplemental Figure 3. Confirmation of Gpc6 ISH probe specificity.**

Schematics of the Gpc6 cDNA highlighting the positions of the ISH probes. Grey lines - UTRs, blue line - coding sequence. Both probes produced similar signal (compare to Fig 2**d**, **g** and **k**). Probe 1 was used in the main study (Fig 2, Supplemental Figure 4). Probe 2 was only used to confirm the specificity of the ISH signal and not in the main study. A sense probe control did not produce any signal.

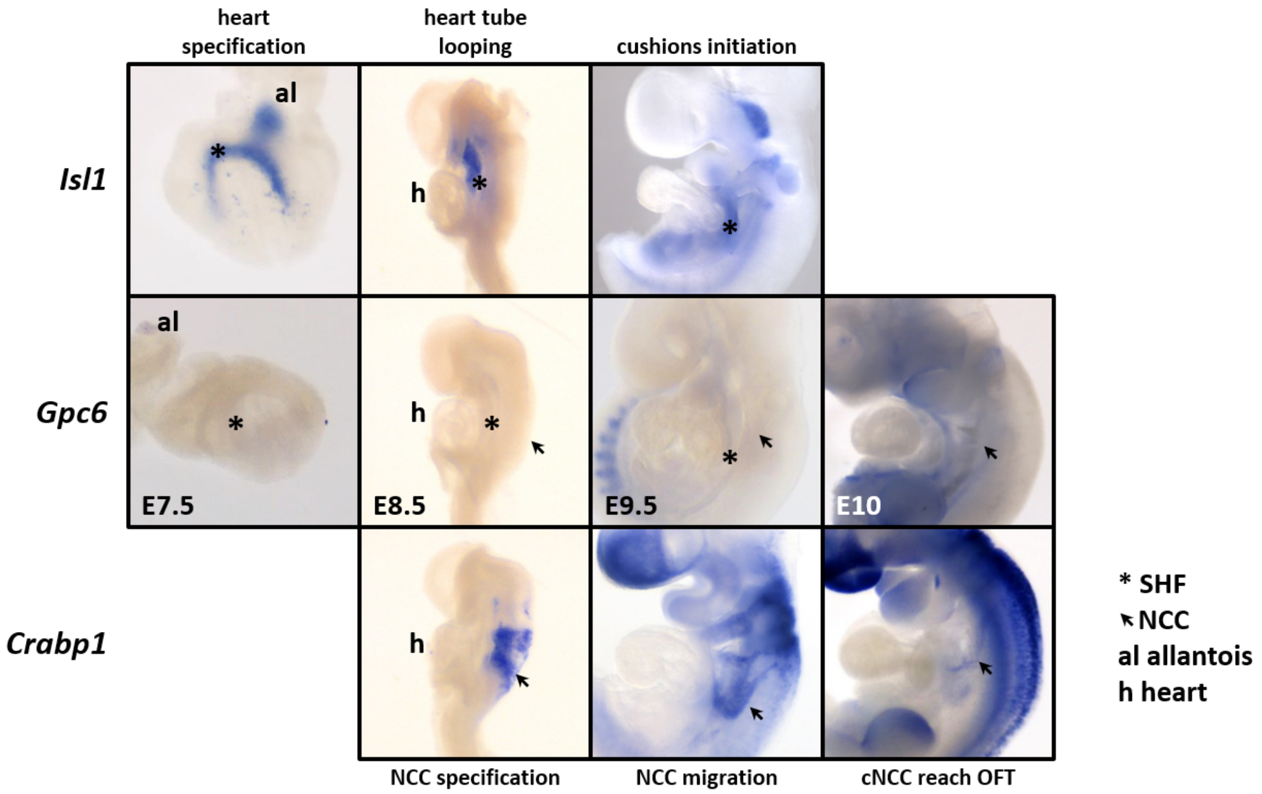


**Supplemental Figure 4. Gpc6 expression during cardiac specification and in migrating NCC.**

*Gpc6* expression was not detected at cardiac mesoderm specification stage (E7.5, labelled by *Isl1* expression), cardiac looping stage (E8.5) or during endocardial cushions initiation (E9.5). Gpc6 was also not expressed in neural crest cells (identified by *Crabp1* expression) at NCC specification (E8.5) and migration (E9.5-10) stages. Asterisk - second heart field (SHF), arrow points to NCC, al allantois, h heart.

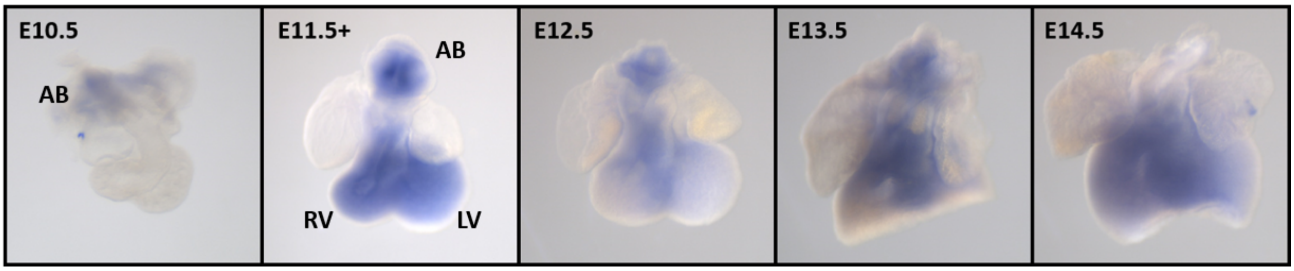


**Supplemental Figure 5. Gpc5 expression.**

Gpc5 expression was assessed in E10.5-14.5 mouse embryonic hearts by ISH. A weak signal was observed in the aortic bulb at E10.5 (AB); at E11.5, the expression spread to the whole OFT and ventricles (RV, LV), where it was further observed in E12.5 through to E14.5.

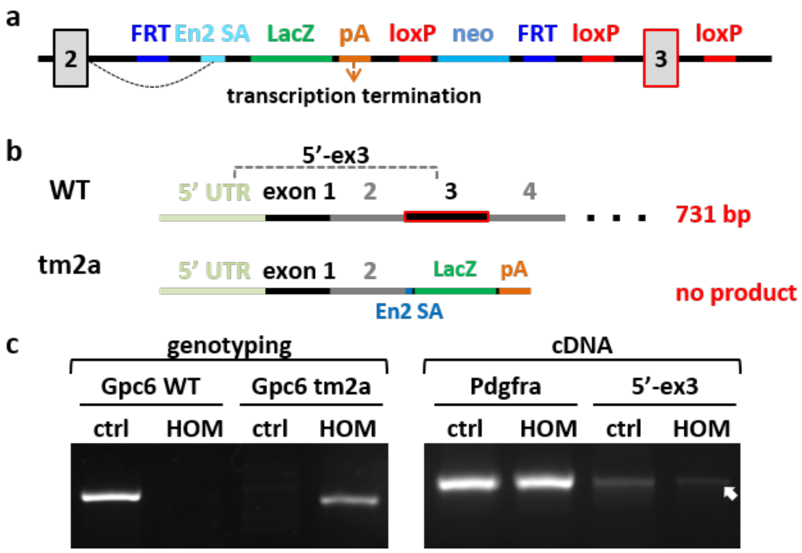


**Supplemental Figure 6. Gpc6 tm2a allele analysis.**

**a** Schematics of the Gpc6$^{tm2a}$ allele creating floxed exon 3 (highlighted with red frame). FRT flip recombinase sites, loxP Cre recombinase sites, En2 SA engrailed-2 splice acceptor site, lacZ β-galactosidase sequence, neo neomycin resistance gene, pA transcription termination signal. **b** Transcripts produced from the WT and tm2a alleles of the Gpc6 gene, and location of the primers (5'-ex3) to detect it. **c** Genotyping PCR confirming the Gpc6 WT or homozygous tm2a alleles (left), expression of *Pdgfra* (as a positive control for cDNA quality) and *Gpc6* (5'-ex3) (right; see genotyping primer schematics in Supplemental Figure 7). Arrow points to the PCR product produced from full-length Gpc6 cDNA which is only expected in control sample. The product was isolated from the gel and sequenced to confirm its identity.

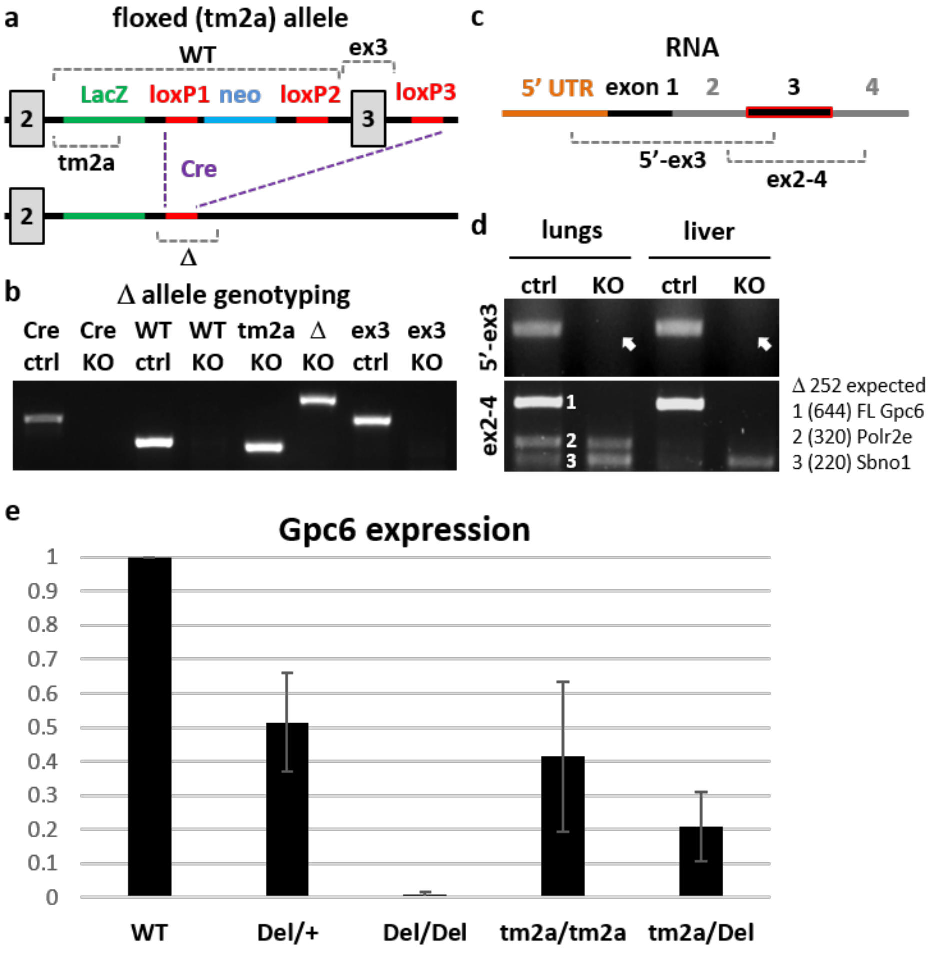


**Supplemental Figure 7. Generation and confirmation of the global Gpc6 knock-out mice.**

After demonstrating that homozygous mice still express full-length *Gpc6* RNA, we generated the global Gpc6 deletion by crossing Gpc6$^{tm2a}$ males to Keratin14-Cre females, which express Cre recombinase protein in oocytes. After the confirmation of the successful KO, further outcrossing removed the K14Cre transgene from the KO animals to avoid undesired effects of K14Cre transgene on phenotype (via disruption of an endogenous mouse gene (Dennis et al. 2012) or Cre toxicity (Pomplun et al. 2007)). **a** Schematics of the floxed Gpc6 allele and generation of the gene knock-out by deletion of the floxed exon 3, leading to a frame shift in protein amino acid sequence. LacZ β-galactosidase gene, loxP Cre recombinase sites, neo neomycin resistance gene, Cre recombinase. Grey dashed lines indicate the location of primers used in genotyping: WT (detects WT Gpc6 allele), tm2a (original tm2a allele), ex3 (presence of intact exon 3) and Δ (presence of the genomic fragment with deleted exon 3). **b** Confirmation of the successful generation of homozygous KO mice by PCR. Animals were genotyped for WT Gpc6 allele (absent in KO tissues), tm2a allele and exon 3 (absent in KO) and Δ fragment with deleted region between loxP sites 1 and 3 containing Neomycin cassette and exon 3 (see Sup Table 1 for primer sequences and expected product sizes). **c** Schematics showing the location of the primers used to detect Gpc6 RNA in control and KO tissues. **d** Gpc6 RNA is no longer detectable on KO tissues (lungs and liver). 5'-ex3 product is missing in KO samples (arrow) while still present in controls. ex2-4 product (expected size 644 bp) amplified from full length Gpc6 RNA was detected in control tissues (1) but absent in KO (arrow). Smaller products (2, 3) were sequenced and confirmed to be non-specifically amplified from different genes (*Polr2a* and *Sbno1*) but not from Gpc6 RNA (expected size of the ex2-4 product for the Δ allele is 252 bp). **e** Quantification of the Gpc6 transcript in the E11.5+ OFT of different genotypes.

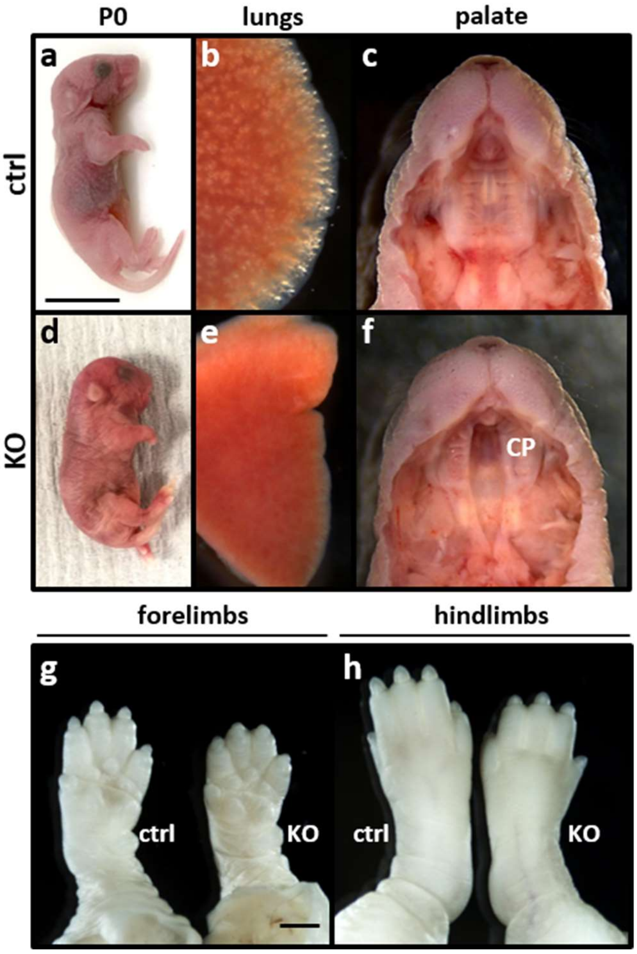


**Supplemental Figure 8. Extra-embryonic defects in Gpc6 KO pups.**

KO pups exhibit shorter bodies (**a** vs **d,** scale bar 1 cm), no air in lungs (**b** vs **e**), and cleft palate (CP, **c** vs **f**). In KO pups, both fore- and hindlimbs were found to be shorter than in control pups. Reduction of size (both limbs and whole body) was clearly noticeable around E18.5. Scale bar 0.1 mm.

**a** **Pathway analysis**

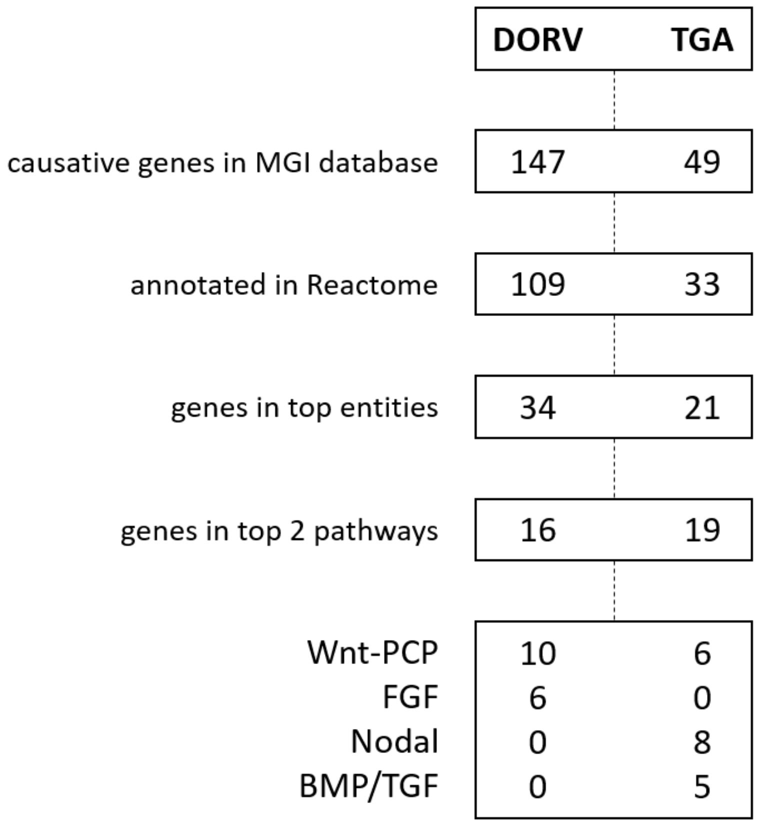


**b** **DORV and TGA overlap**

| | |
|---|---|
| Cfc1 | Nodal |
| Foxh1 | Nodal |
| Lefty1 | Nodal |
| Lefty2 | Nodal |
| Zic3 | Plur SC |
| Pitx2 | TFAP-2 |
| Dvl2 | Wnt |
| Prickle1 | Wnt |
| Psme4 | Wnt, Mapk |

**Supplemental Figure 9. Signalling pathway analysis.**

**a** DORV and TGA were selected as primary phenotypes present in the Gpc6 KO hearts. Causative genes for both pathways (in mouse model) were identified in the analysis of MGI database and run through the Reactome algorithm. Identified entities were grouped by the pathway they represent, and found genes counted. **b** The genetic overlap between DORV and TGA phenotypes. The genes identified in "genes in top entities" were compared and common genes are shown, with the corresponding pathway.

**Supplemental Table 1. List of primers used in current study.**

MC mesenchymal cells, SHF second heart field. Underlined is the site of T7 (in For primer) or T3 (in Rev primer) polymerase binding.

| aim | primers set | For primer | Rev primer | product size |
|---|---|---|---|---|
| GPC6-AS2 detection | GENECODE annotation set 1 | ACATTTAGGTTCAAAGCCAGGAG | TGCACACACGTAAACTTTCAAA | 212 RNA<br>613 genomic |
| | GENECODE annotation set 2 | GTGGCAAAGCACCTGTTGTC | TGATAAATGCACACACGTAAACT | 194 RNA<br>595 genomic |
| | RefSeq annotation set 3 | CCCCTTCTCTGTGGCATTGA | TGCACACACGTAAACTTTCAAA | 274 RNA |
| | RefSeq annotation set 4 | GGAATAGCCCCTTCTCTGTGG | TGATAAATGCACACACGTAAACT | 288 RNA |
| GPC6 detection | mouse *Gpc6* ISH 1 | TAATACGACTCACTATAGGGTTG<br>GCTTGCCATCCTCCATC | AATTAACCCTCACTAAAGGGCCG<br>AGCCCAGAAGTCATTGA | 751 For-T7<br>731 For only |
| | mouse *Gpc6* ISH 2 | AGCGATGAATCCAGTGGCTC | AATTAACCCTCACTAAAGGGGCG<br>CTTTGTAAACCTCGCTT | 758 |
| | human *GPC6* | ACACGTTTCAGGCCCTACAA | TGTGGCATGGTCCATTGTCC | 679 |
| cushions MC marker | *Pdgfra* | TGGAAGTGGTTAACGCCTCC | AATTAACCCTCACTAAAGGGCAC<br>CTCCACCACGAACTCTC | 784 |
| SHF marker | *Isl1* | TTTCCCTGTGTGTTGGTTGC | AATTAACCCTCACTAAAGGGCTC<br>AGTACTTTCCAGGGCGG | 864 |
| Gpc6 genotyping | WT | GCTATCCAATGGCCCCTTTG | ATTCTTCGTTCCACCTGAGC | 531 |
| | tm2a | CCCCTTTCCAAACACAATCAG | CCCTCTCACCTTCTACCCCA | 481 |
| | $\Delta$ | TCCTGGAGCCCGTCAGTATC | TGCAGCGCTGACCTCATAAA | 905 |
| | exon 3 | GCCTACCATATGCGAGGACC | CGTATGGAATGGCTTGGCAT | 714 |
| Gpc6 cDNA | exon 2-4 | GGAATATACCTGCTGCACCAC | AGCCCTTCATGACGTTGAGG | 644 FL<br>252 $\Delta$ |
| Cre transgene | Cre | GCCAGCTAAACATGCTTCATC | ATTGCCCCTGTTTCACTATCC | 727 |
| qPCR primers | *Ki67* | ATCATTGACCGCTCCTTTAGGT | GCTCGCCTTGATGGTTCCT | 104 |
| | *Tpx2* | GATGCCCCCACCGACTTTATC | CTTGTTCTCCAAGTTGGCCTT | 102 |
| | *Top2A* | CAACTGGAACATATACTGCTCCG | GGGTCCCTTTGTTTGTTATCAGC | 185 |
| | *Bcl2* | GTCGCTACCGTCGTGACTTC | CAGACATGCACCTACCCAGC | 284 |
| | *Casp2* | TACTCCCACCGTTGAGCTGT | CCGTAGCATCTGTGGATAGGC | 102 |

| | | | | |
|---|---|---|---|---|
| | *Casp8* | TGCTTGGACTACATCCCACAC | TGCAGTCTAGGAAGTTGACCA | 169 |
| | *Adam19* | GTCCTTTCTTAGTTGGAGGCG | CATGCTAACTCCTCCAGACTG | 151 |
| | *Pdgfra* | CGTTGACCTGCAGTGGACTT | GAGTTTGATCTCCTCCAGCATG | 73 |
| | *Pdgfrb* | TGCTGGGAAGAAAAGTTTGAGA | GGCTTGGGACCTCAGGAT | 153 |
| | *Rxra* | AAGACCTGACCTACACCTGC | GTTCTCATTCCGGTCCTTGC | 161 |
| | *Snai1* | CGGAAGCCCAACTATAGCGA | GTCGTAGGGCTGCTGGAAG | 66 |
| | *Snai2* | AGCGAACTGGACACACACA | TGCTCCCGAGGTGAGGATC | 99 |
| | *Tbx20* | CACGGCCTCCTTGCTCAATC | GAGTCCCGGAATCCTTTGGC | 150 |
| | *Twist1* | GTCCCACTAGCAGCGGAG | AGACTGTCCATTTTCTCCTTCTC | 88 |
| | *Cnn2* | CCAAAAGGAAGCAGAACTCCG | GAGCGGTTGATTTTAGGGACG | 150 |
| | *Daam1* | CCTCACAGACAAACACAGGG | AGCAGCCATGGAATTGAGCT | 151 |
| | *Dvl1* | TACCCCTACCAGTACCCAGG | TGGTCTGATTCACTGCCACT | 203 |
| | *Ptk7* | CGCTGAGAGTGATACTGGTG | AGATCACAGTGGGCTTGGG | 191 |
| | *Vangl2* | AATCAGTGACGATCCAGGCT | ACACTGTCCTCCATGTCCTT | 172 |
| | *Ipo8 (internal control)* | AACAGACCCGAACTTTGACC | GATTCTGCAGGAACAGCTCA | 122 |

**Supplemental Table 2. Neonatal lethality of Gpc6 KO pups.**

Assessment of the viability of the Gpc6 KO animals. E10.5-16.5 embryos and 3 weeks old mice were genotyped.

| | | +/+ | Δ/+ | Δ/Δ | P value ($\chi^2$ test) |
|---|---|---|---|---|---|
| embryos (n=105) | observed | 24 | 46 | 35 | 0.3345 |
| | expected | 26.25 | 52.5 | 26.25 | |
| adults (n=103) | observed | 47 | 56 | 0 | <0.0001 |
| | expected | 25.75 | 51.5 | 25.75 | |

**Sup Table 3. Viability of the tissue-specific KO mice.** Intercross Cre:Gpc6$^{tm2a/+}$ X Gpc6$^{tm2a/tm2a}$.

| | | tm2a/+ | tm2a/tm2a | Cre tm2a/+ | Cre tm2a/tm2a | P value ($\chi^2$ test) |
|---|---|---|---|---|---|---|
| **Tie2Cre** (n=99) | observed | 24 | 19 | 29 | 27 | 0.5139 |
| | expected | 24.75 | 24.75 | 24.75 | 24.75 | |
| **Wnt1Cre** (n=69) | observed | 27 | 19 | 23 | 0 | <0.0001 |
| | expected | 17.25 | 17.25 | 17.25 | 17.25 | |

**Sup Table 4. The summary of lethality and defects observed in Gpc6 conditional and global KO strains.**

OAo overriding aorta, AoS aortic stenosis, PS pulmonic stenosis, VSD ventricular septal defect, DORV double outlet right ventricle, r-Ao rightward mal-positioned aorta, RVH right ventricular hypertrophy.

| KO | lethality | lungs | palate | heart defects | penetrance |
|---|---|---|---|---|---|
| Tie2Cre (endoMES) | viable | normal | normal | no defects | - |
| Wnt1Cre (cNCC) | neonatal | no air | cleft | no defects | - |
| double KO | neonatal | no air | cleft | OAo + VSD | 75% |
| global KO | neonatal | no air | cleft | DORV + r-Ao + VSD<br>AoS, PS, RVH | 100% |
